# Centrosomes act as organizing centers to promote Polo kinase-mediated adaptation to persistent DNA damage

**DOI:** 10.1101/2024.06.13.598907

**Authors:** Laurence Langlois-Lemay, Damien D’Amours

**Affiliations:** Ottawa Institute of Systems Biology, Department of Cellular and Molecular Medicine, University of Ottawa, Roger Guindon Hall, 451 Smyth Rd, Ottawa, ON, K1H 8M5, Canada

**Keywords:** Cdc5, PLK1, MTOCs, centrosomes, spindle pole bodies, cell cycle, adaptation

## Abstract

The ability of cells to overcome cell cycle arrest and adapt to the presence of unrepairable DNA damage is under the control of polo-like kinases (PLKs) in eukaryotes. How DNA damage checkpoints are silenced or bypassed during the adaptation response is unknown, but the process requires enrichment of the Cdc5 PLK to microtubule organizing centers (MTOCs), such as the yeast centrosomes or spindle pole bodies (SPBs). Here we found that SPBs play an active role as supramolecular organizing centers that coordinate Cdc5 recruitment and signaling to downstream effectors during the adaptation response to DNA damage. We show that SPB components Nud1, Spc110, and Spc72 are key effectors of Cdc5 recruitment to SPB in the presence of sustained DNA damage. Following recruitment, Cdc5 transduces a phospho-signal to key structural subunits of the SPB, including Cnm67 and Mps3. We demonstrate these phosphorylation events are required to bypass cell cycle checkpoint arrest and enable effective adaptation to DNA damage. This response is specific because it cannot be recapitulated by a generic inactivation of MTOC activity. Collectively, our results indicate that centrosomes can act as supramolecular platforms to coordinate dynamic recruitment and substrate selection of PLKs during the DNA damage response.

## Introduction

The maintenance of genomic integrity is an essential aspect of cell survival and homeostasis for all living species. In eukaryotic cells, the DNA damage response (DDR) integrates a highly conserved set of signaling pathways established to safeguard genomic integrity in the face of genotoxic stress (reviewed in Paulovich et al., 1997; Pizzul et al., 2022). Detection of DNA lesions in damaged cells induces a reversible checkpoint-mediated cell cycle arrest that allows time for repair of DNA lesions prior to DNA synthesis and/or mitotic re-entry (Weinert and Hartwell, 1988). Through a process termed recovery, cells that successfully complete DNA repair return to active proliferation by deactivation of checkpoint effectors (reviewed in Yam et al., 2022). Cells incapable of repairing DNA lesions will engage in apoptosis when they carry high levels of persistent DNA damage, a behavior that protects organisms from genomic instability and oncogenesis. Interestingly, cells experiencing low or moderate levels of unrepairable DNA damage can bypass checkpoint-mediated cell cycle arrest and evade apoptosis through a specific process known as the adaptation response to persistent DNA damage. By turning off the G2/M checkpoint, cells can ignore persistent DNA damage and resume cycling despite a compromised genome (Sandell and Zakian, 1993; Toczyski et al., 1997; Lee et al., 1998b). This evolutionarily conserved pathway can maximize cell survival by making possible the repair of lingering DNA lesions in subsequent rounds of cell division (Galgoczy and Toczyski, 2001), and/or by allowing asymmetric partition of damaged DNA to only one of the two daughter cells after mitosis (Kaye et al., 2004; reviewed in Serrano and D’Amours, 2014). To be successful, this adaptation response requires a careful coordination of checkpoint deactivation mechanisms with cell cycle re-entry. This coordination is controlled by polo-like kinases (PLKs) in all eukaryotes (reviewed in Serrano and D’Amours, 2014; Langlois-Lemay and D’Amours, 2022).

PLKs are conserved serine/threonine protein kinases involved in the regulation of cell division and the maintenance of genomic integrity (Llamazares et al., 1991; Kitada et al., 1993; Golsteyn et al., 1994). Among the five PLKs found in humans (PLK1-PLK5), PLK1 is the most central in events related to cell cycle progression (Kalous and Aleshkina, 2023). The key structural features of PLKs include an N-terminal kinase domain (KD) connected to a C-terminal phospho-serine/threonine binding module known as the Polo-box domain (PBD). This family of kinase typically phosphorylates its targets via a two-stage process. First, the PBD of PLKs acts as a signal recognition module to identify and bind cellular targets that have been pre-phosphorylated by CDK1 *in vivo* (Lee et al., 1998a; Elia et al., 2003a; Elia et al., 2003b). Second, the PBD-induced proximity of PLK targets to the KD of the enzyme leads to their hyperphosphorylation and amplification of the signal initiated by CDK1 phosphorylation. Ultimately, this cascade of events promotes phospho-signal transduction to stimulate downstream events critical for cell cycle progression (reviewed in Kalous and Aleshkina, 2023).

Human PLK1 and its budding yeast homolog, Cdc5, are highly dynamic proteins that must enrich at various subcellular locations, such as the centrosome, to promote cell cycle progression (Song et al., 2000; Seong et al., 2002). The budding yeast spindle pole body (SPB) is analogous to the human centrosome in that it functions as the main microtubule-organizing center (MTOC) and represents an important scaffolding site for the enrichment of Cdc5 kinase. While best known for its MTOC function, the centrosome also plays an essential role as a supramolecular platform on which several kinases enrich to mediate key signaling events (reviewed in Langlois-Lemay and D’Amours, 2022). Consistent with this notion, the PBD-mediated localization of Cdc5 at SPBs is crucial for mitotic exit, cytokinesis, maintenance of SPB integrity, and ploidy control (reviewed in Botchkarev and Haber, 2018). Moreover, recruitment of Cdc5 to SPBs is also essential for DNA damage adaptation and directly impacts cell survival (Ratsima et al., 2016; Rawal et al., 2016). A Cdc5 mutant defective for PBD-mediated substrate recognition does not localize to the SPB and fails to undergo DNA damage adaptation (Ratsima et al., 2011; Ratsima et al., 2016). The PBD-mediated localization of Cdc5 to SPBs is therefore essential to modulate cell proliferation upon checkpoint activation and can directly impact cell homeostasis.

In this study, we explore the specific contribution of SPBs in Cdc5-dependent adaptation to DNA damage. We took advantage of the bipartite structure of PLKs to reveal the identity of two separate pools of effectors of Cdc5 at the SPB and found that several SPB constituents are direct targets for this kinase in damaged cells. Our work sheds light on the choreography of interactions connecting Cdc5 to diverse SPB components during adaptation to DNA damage and reveals the specific sequence of events required to bypass checkpoint activation and return damaged cells to a cycling state.

## Results

### Identification of candidate receptors for Cdc5 at the SPB

We previously showed that PBD-mediated binding of Cdc5 to SPBs is essential to promote adaptation to DNA damage (Ratsima et al., 2016). To identify the putative SPB receptor protein(s) for Cdc5, we examined the sequence of SPB components for the presence of the conserved phospho-serine/threonine-binding motif recognized by Cdc5 PBD (i.e., S-[pS/pT]-[P/X]; Elia et al., 2003a; Elia et al., 2003b). **Figure 1A** shows the number of PBD-binding motifs identified in each of the 17 components of the yeast SPB. This analysis revealed Nud1 as a top receptor candidate for Cdc5 because it contains the highest number of PBD-binding consensus sites (15) among all SPB components.

**Figure 1.**
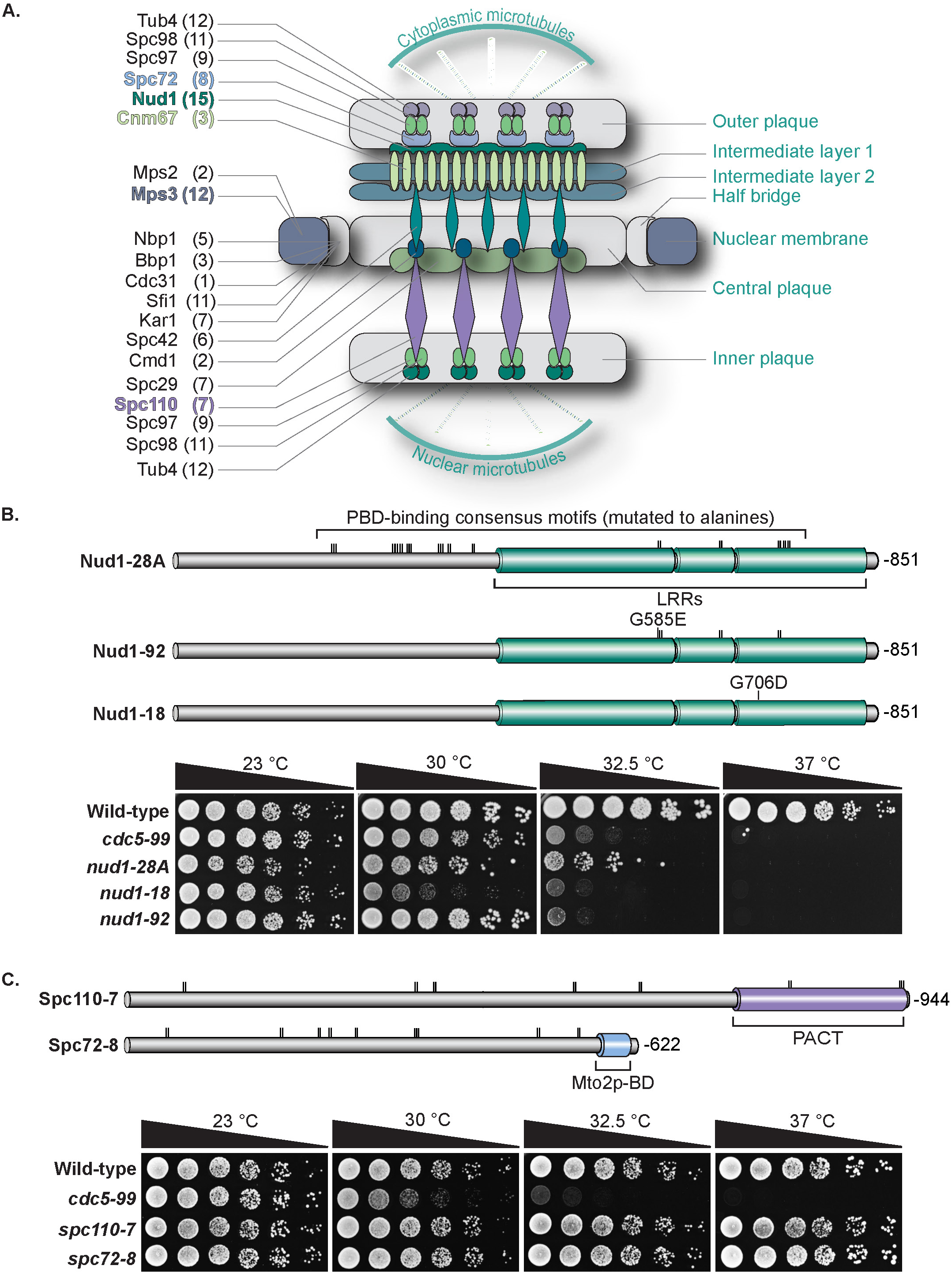
Growth phenotype of novel SPB mutants. **(A)** Schematic representation of the yeast SPB and its physical components. Number of consensus binding sites for Cdc5 PBD are reported for each SPB protein (see number in brackets). Subunit arrangement within the SPB is derived from Klenchin (2011), Jaspersen and Winey (2004), and Kilmartin (2014). **(B)** Schematic representation of Nud1 protein domains and mutations (top). The linear domain architecture and location of mutations (vertical lines) are shown to scale. Protein length is shown on the right side of each structure. LRRs: Leucine-rich repeats. All mutations are serine/threonine to alanine unless stated otherwise. The proliferation phenotype of *nud1* mutants at various temperatures is shown under the protein schematics. Cells were spotted on YPD media and grown at the indicated temperatures. (n=3) **(C)** Domain architecture and position of mutations inserted in Spc110-7 and Spc72-8 proteins (top). Representation of domains and mutations are as described in panel B. PACT: Pericentrin-AKAP450 centrosomal targeting domain. Mto2p-BD: Mto2p-binding domain. Growth properties of *spc110-7* and *spc72-8* mutants (bottom) were assessed on rich YPD media at 23 °C, 30 °C, 32.5 °C and 37 °C. (n=3)

To assess the role of Nud1 as a SPB recruitment factor for Cdc5 during adaptation to DNA damage, we created three *nud1* temperature-sensitive (ts) alleles of this gene. The first allele, *nud1-28*, has lost all putative Cdc5 PBD-binding sites in *NUD1* (**Figure 1B** – top). The second allele, *nud1-92*, is a milder version of the *nud1-28* mutant and contains only six PBD-binding mutations along with a G585E substitution (**Figure 1B** – middle schematic; Alexandar et al., 2004). Finally, the third *nud1* allele corresponds to the *cdc18-1* mutation originally described by Hartwell and colleagues (Hartwell et al., 1973; Bardin and Amon, 2001) and carries a G706D point substitution (**Figure 1B** – bottom schematic). For clarity, the *cdc18-1* allele will be referred to as *nud1-18*. Consistent with the essential nature of *NUD1*, all three alleles we tested conferred varying degrees of temperature-sensitivity to yeast in serial dilution growth assays, with the *nud1-18* mutant being the most affected at high temperature (**Figure 1B**).

Aside from Nud1, two core SPB components –namely Spc110 and Spc72– have been reported to show affinity for Cdc5 (Park et al., 2004; Snead et al., 2007), making them plausible candidate receptors for Cdc5 during adaptation to DNA damage. Spc110 and Spc72 contain seven and eight PBD-binding motifs (**Figure 1A)**, respectively, and can bind to Cdc5 PBD *in vitro* (Snead et al., 2007). To evaluate their contribution as SPB recruitment factors for Cdc5, we constructed the *spc110-7* and *spc72-8* alleles in which all the consensus PBD-binding motifs were rendered inactive by mutation of the putative phospho-serine/threonine residues to alanines (**Figure 1C** – top). Neither *spc110-7* nor *spc72-8* mutants showed proliferation defects at 37 °C (**Figure 1C** – bottom), indicating that removal of PBD-specific binding sites in these structural components of the SPB does not interfere with the essential/MTOC function of the organelle.

### Nud1 contributes to the adaptation response to persistent DNA damage

Using the mutants described above, we next investigated the contribution of Nud1 to the adaptation response of yeast experiencing unrepairable DNA damage. Persistent DNA damage was generated by temperature-induced (31–33 °C) inactivation of the *cdc13-1* allele, resulting in a sustained DNA damage response and G2/M checkpoint-mediated cell cycle arrest (Weinert et al., 1994). The duration of this DNA damage-induced cell cycle arrest normally ranges from 8 to 10 hours, after which cells adapt to the presence of DNA damage and re-enter a proliferative state (Toczyski et al., 1997). As previously observed, *cdc13-1* mutant cells showed a typical large-budded morphology 2.5 hours after exposure to 32.5 °C, reflecting a robust G2/M checkpoint arrest. These cells eventually form microcolonies comprising at least 5 cell bodies of normal size 24 hours after formation of DNA lesions, consistent with an effective adaptation response to DNA damage. In contrast, *nud1* mutants showed a strongly reduced microcolony formation capacity and a steady increase in cell size over the course of the experiment. This morphological feature, also reported in the adaptation mutant *cdc5-16* (Ratsima et al., 2016), represents a consequence of an active metabolism in non-dividing cells (Johnston et al., 1977). Adaptation rates in *nud1* mutants only reached 30% to 50% of the rate observed in a *NUD1 cdc13-1* control strain (**Figure 2B**). Similar results were obtained when *nud1* mutants were tested for adaptation in response to a sustained HO endonuclease-induced double-strand break (DSB) (**Figure S1**). Thus, Nud1 contributes to the adaptation response to DNA damage and its effects are observed with different alleles and forms of DNA damage.

**Figure 2.**
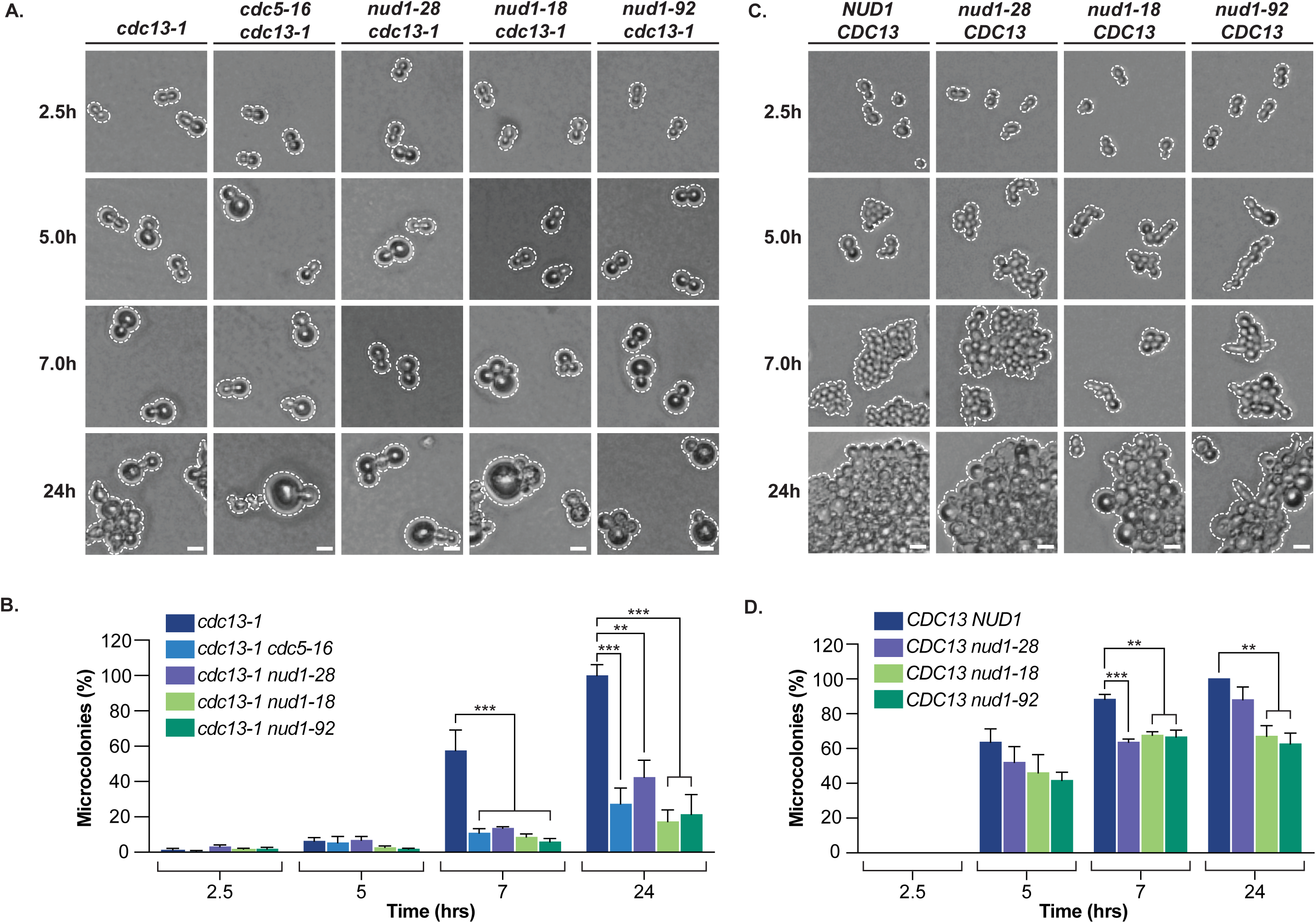
Nud1 is required for the adaptation response to persistent DNA damage. **(A)** *nud1* mutants were assayed for microcolony formation after inactivation of *cdc13-1* at restrictive temperature (32.5 °C) and formation of unrepairable DNA damage at telomeres. All *nud1* mutants showed reduced rates of adaptation to DNA damage relative to the control *NUD1* strain. Scale bars correspond to 10 µm. Outlines of cells and microcolonies are marked with dashed lines to facilitate visualization. Strain genotypes are described in the panel headings. **(B)** Quantification of the fraction of individual cells growing into microcolonies for each timepoint tested in panel A. A minimum of 100 cells per strain was counted for each timepoint. All values were normalized to 100% based on the T24 h mean value of the *cdc13-1* control strain. Data represented as normalized mean values and SEM of at least 3 independent experiments. Statistical analysis performed using a one-way ANOVA test on normally distributed values with Dunnett’s multiple comparisons post hoc test. Statistical significance is shown on graph with asterisks (** is p ≤ 0.01; *** is p ≤ 0.001). **(C)** Microcolony formation assays performed with *nud1* mutants in the absence of DNA damage (i.e., wild-type *CDC13* background at 32.5 °C). Scale bars correspond to 10 µm. **(D)** Quantification of *CDC13* cells forming microcolonies at each timepoint shown in panel C. A minimum of 100 cells per strain was counted for each timepoint. Data representation and analysis of statistical significance are as described in panel B.

To verify that the adaptation phenotype detected in *nud1* mutants was specific to the adaptation pathway and did not stem from other proliferation defects, we evaluated the microcolony formation properties of *nud1* strains growing at 32.5 °C in the absence of DNA damage (i.e., in a *CDC13* background). While appearance of microcolonies was slightly delayed in *nud1* mutants compared to wild-type cells, we observed that *nud1* mutants could form microcolonies at rates ranging from 70%–90% of wild-type rates under non-damaging conditions (**Figure 2D**). Importantly, strains carrying the *nud1-28* allele showed only partial temperature sensitivity for growth at 32.5 °C (**Figure 1B**) and could form microcolonies at nearly wild-type frequency in the absence of DNA damage (**Figure 2D**), despite showing a strong adaptation defect in the presence of DNA damage (i.e., with *cdc13-1*; **Figure 2B**). Therefore, the microcolony formation defect of the *nud1* alleles we tested mostly reflects the role of Nud1 protein in the adaptation response to DNA damage.

### Nud1 operates upstream of Cdc5 during the DNA damage response

To understand how Nud1 contributes to Cdc5-mediated adaptation to DNA damage, we next aimed to determine the relative position of Nud1 in this pathway. To achieve this, we used *CDC5* overexpression as a tool to place Nud1 upstream or downstream of Cdc5 point-of-action, as previously performed by us and others (Ratsima et al., 2016; Vidanes et al., 2010). Wild-type *CDC5* was overexpressed under the *GAL1* promoter in *cdc13-1* cells defective for *nud1* and the kinetics of adaptation were monitored in the presence of DNA damage. As seen in **Figure 3A**, Cdc5 overexpression allowed for markedly increased adaptation kinetics in *nud1* mutants relative to non-overexpressing cells (c.f., **Figure 2B**). *nud1* mutants overexpressing *CDC5* showed an abridged G2/M cell cycle arrest and adaptation rates mirroring or exceeding those observed in *NUD1* control cells 24 hours after induction of DNA damage (**Figure 3B**). The suppression of the *nud1* adaptation defect by *CDC5* overexpression indicates that this protein operates upstream of (or in parallel to) Cdc5 during the cellular response to DNA damage.

**Figure 3.**
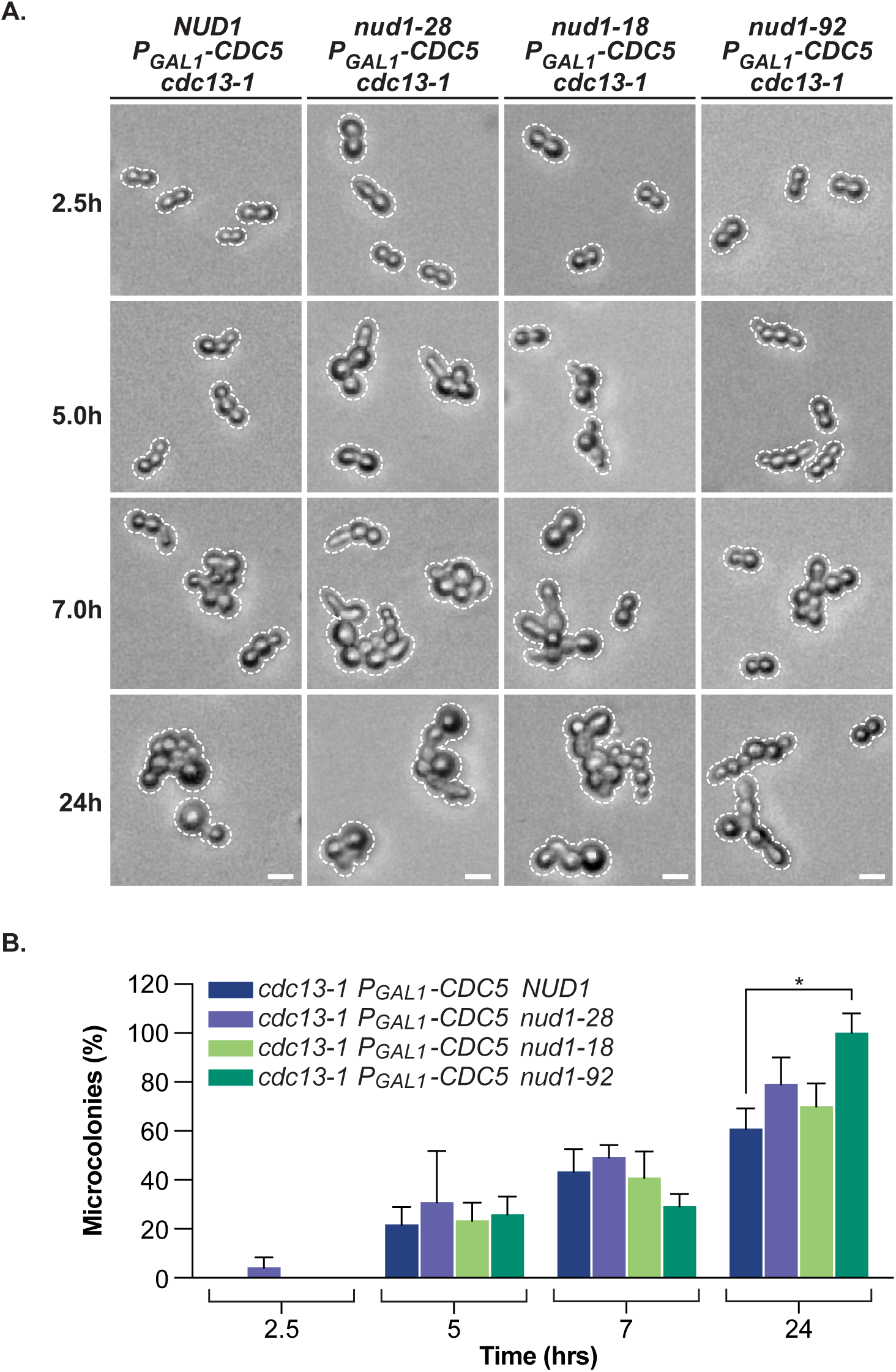
Nud1 operates upstream of Cdc5 in the adaptation response to DNA damage. **(A)** *nud1* strains overexpressing *CDC5* (*P_GAL1_-CDC5*) were assayed for microcolony formation in a DNA damage adaptation test using *cdc13-1* inactivation at 32.5 °C. Cells were plated on solid YEP medium containing 2% galactose (YEPG). Overexpression of *CDC5* in *nud1* mutants rescued the adaptation phenotype and led to an abridged G2/M cell cycle arrest. Scale bars, genotype representation and cell outlines are as described in Figure 2. **(B)** Quantification of the fraction of cells growing into microcolonies for each strain tested in panel A. A minimum of 100 cells per strain was counted at each timepoint. All values were normalized to 100% based on the T24 h mean value of the *P_GAL1_-CDC5 cdc13-1 nud1-92* mutant strain, as it presented the highest adaptation rates observed in this experimental dataset. Data represented as mean values and SEM of at least 3 independent experiments. Statistical analysis performed using a one-way ANOVA test on normally distributed values with Dunnett’s multiple comparisons post hoc test. Statistical significance is shown on graph with asterisks (* is p ≤ 0.05).

### Spc110 and Spc72 assist Nud1 in Cdc5-dependent adaptation to DNA damage

Previous reports have shown that Cdc5 PBD can bind *in vitro* to other SPB components aside from Nud1 (Park et al., 2004; Snead et al., 2007). In particular, SPB inner and outer plaque components Spc110 and Spc72 are promising candidates as SPB receptors for Cdc5 because they contain multiple consensus sites for PBD binding (**Figure 1A**). To test this hypothesis, we designed *spc110-7* and *spc72-8* mutants in which all putative Cdc5 phospho-binding sites were mutated from serines/threonines to alanines (see **Figure 1C**). Testing these mutants for adaptation to DNA damage showed that neither *spc110-7* nor *spc72-8* are defective in microcolony formation following a G2/M checkpoint arrest (**Figure S2**). As previous literature highlighted the preferential binding of Cdc5 PBD to Nud1 at the SPB *in vitro* (Park et al., 2004), we projected that the contribution of other SPB components as Cdc5 recruitment factors in adaptation might be better assessed in a Nud1-defective background. Interestingly, double mutants carrying *nud1-28* and *spc72-8* alleles could not be obtained due to systematic reversion of *spc72-8* to wild-type following sporulation of heterozygous diploids. Instead, we used a milder PBD-defective allele, *nud1-92*, to create a triple mutant with *spc110-7* and *spc72-8*. Growth assays on solid medium indicated that *spc110-7* and *spc72-8* single mutants proliferate akin to wild-type control at all temperatures tested (cf., **Figure 1C**). In contrast, combining these alleles with *nud1-92* led to severely impaired growth, even at room temperature (23 °C) (**Figure 4A**), reflecting a clear synthetic effect associated with the combination of PBD mutations in several SPB components.

**Figure 4.**
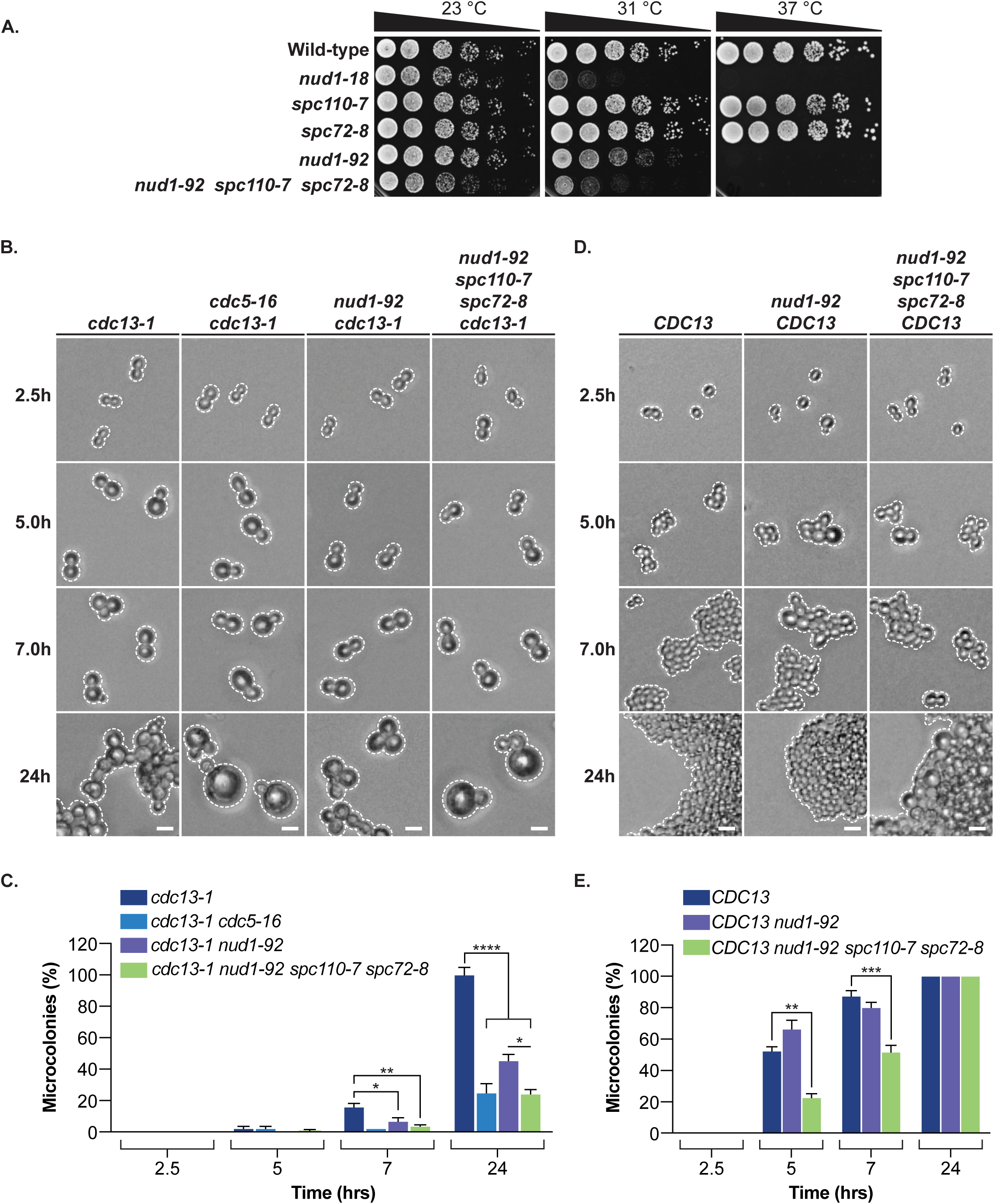
Spc110 and Spc72 contribute to Cdc5-dependent adaptation to DNA damage. **(A)** Proliferation properties of *spc110-*7, *spc72-8*, and *nud1-92* mutants at various temperatures. Cells were spotted on the surface of solid YPD medium and grown at 23 °C, 31 °C and 37 °C. The combination of the three SPB alleles into a triple-mutant exacerbated the growth phenotype of *nud1-92*. (n=3) **(B)** Impact of *spc110-*7 and *spc72-8* on the adaptation defect of *nud1-92*. Microcolony formation following DNA damage induction was assessed at 31 °C, a restrictive temperature for *cdc13-1* (Garvik et al., 1995). The adaptation defect of the triple-mutant was twice as severe relative to *nud1-92* and recapitulated the adaptation phenotype of *cdc5-16* PBD-binding mutant. Scale bars, genotype representation and cell outlines are as described in Figure 2. **(C)** Quantification of individual cells growing into microcolonies for each timepoint in controls, *nud1-92* and triple mutant strains in the adaptation test shown in panel B. A minimum of 100 cells per strain was counted for each timepoint. Data represented as normalized mean values and SEM of at least 3 independent experiments. Statistical analysis performed using a one-way ANOVA test on normally distributed values with Dunnett’s multiple comparisons post hoc test. An unpaired t-test on normally distributed values was used to assess the statistical significance of differences observed between *nud1-92* and *nud1-92 spc110-7 spc72-8* strains at T24 h. Statistical significance is depicted on the graph with asterisks (* is p ≤ 0.05; ** is p ≤ 0.01; **** is p ≤ 0.0001). **(D)** Microcolony formation assay performed at 31 °C with control cells, *nud1-92* single mutant and SPB triple-mutant in the absence of DNA damage (*CDC13*). **(E)** Quantification of microcolony formation phenotypes shown in panel D. A minimum of 100 cells per strain was counted for each timepoint. Data represented as mean values and SEM of at least 3 independent experiments. Statistical analysis performed using a one-way ANOVA test on normally distributed values with Dunnett’s multiple comparisons post hoc test. Statistical significance is shown on graph with asterisks (** is p ≤ 0.01; *** is p ≤ 0.001).

Next, we assessed the adaptation behavior of the *nud1-92 spc110-7 spc72-8* triple mutant strain. This experiment was carried out at a lower restrictive temperature of 31 °C to account for the enhanced temperature sensitivity of the triple mutant and to widen the scope of detectable phenotypic variations between *nud1-92* and the triple-mutant. The microcolony assay depicted in **Figure 4** shows that a combination of PBD-binding mutations (i.e., *nud1-92 spc110-7 spc72-8)* led to adaptation defects twice as severe as those observed in the *nud1-92* single mutant and recapitulated the adaptation phenotype observed in *cdc5-16* cells with a 70% decrease in adaptation rates relative to *cdc13-1* control (**Figure 4C**). To validate that this adaptation phenotype was unrelated to impaired cell cycle progression, we measured dynamics of microcolony formation at 31 °C in absence of DNA damage *(CDC13)* in wild-type control cells, *nud1-92* mutant and triple SPB mutant. **Figure 4D-E** shows that while the triple mutant proliferated with slightly slower kinetics relative to wild-type control, the ability of this strain to undergo cell division was maintained in the absence of DNA damage, and growth rates reached *CDC13* wild-type levels 24 hours post temperature shift. As the combination of PBD-binding mutations into the triple mutant exacerbated *nud1-92* adaptation phenotype without impeding other cell cycle processes, we conclude that Nud1, Spc110, and Spc72 are required to produce a full adaptation response to DNA damage.

### Generic inactivation of SPBs does not cause a specific defect in the adaptation response to DNA damage

Given the role played by Nud1, Spc110, and Spc72 in the DNA damage response, we next asked whether generic inactivation of any SPB component would produce equivalent defects in the microcolony formation assay. We introduced the previously described Q110P substitution into Spc42 (Neumüller et al., 2013), a core SPB component involved in SPB duplication (Donaldson and Kilmartin, 1996), to create the *spc42-Q110P* ts mutant (**Figure S3A**). To test whether *spc42-Q110P* inactivation could induce a specific defect in adaptation, we monitored the dynamics of microcolony formation in a *cdc13-1* background at restrictive temperature (32.5 °C). Under these conditions, the *spc42-Q110P* mutant showed a uniform population of large-budded cells arrested in G2/M 2.5 hours after inactivation of *cdc13-1*, consistent with a robust checkpoint-induced arrest, and these cells ultimately formed microcolonies at 40% of the rates observed in a control *SPC42* strain exposed to DNA damage for 24 hours (**Figure S3B-C**). Interestingly, very similar results were observed when the *spc42-Q110P* mutant was tested in the absence of DNA damage (*CDC13*; **Figure S3D-E**). Since the cell cycle arrest and microcolony formation defect occurred regardless of DNA damage in *spc42-Q110P* cells, we conclude that generic inactivation of a SPB core component leads to a loss of MTOC activity that interferes with further cell division. This phenotype contrasts with the ability of *nud1* PBD mutants to form microcolonies in the absence of DNA damage. Taken together, these results indicate that the *cdc13*-induced microcolony phenotype observed with PBD-binding mutants of Nud1 (alone or in combination with *spc110-7* and *spc72-8*) reflects a specific role for this protein in adaptation to DNA damage.

### Cdc5 association with SPB is impaired in adaptation-defective SPB mutants

Since PBD-mediated enrichment of Cdc5 at the SPB is essential to promote checkpoint adaptation (Ratsima et al., 2016) and the SPB triple mutant described above is defective for this pathway, we next asked whether the reduced number of available PBD-binding sites in this strain impairs Cdc5 recruitment at SPBs. To test this possibility, we monitored GFP-Cdc5 localization at the SPB in wild-type, *nud1-92*, and *nud1-92 spc110-7 spc72-8* triple mutant strains expressing the SPB marker Spc29-RFP. We assessed Cdc5-SPB colocalization before DNA damage, in presence of DNA damage, and after recovery from DNA damage by transiently inactivating *cdc13-1* at 31 °C. Whereas control and *nud1-92* strains showed typical SPB behavior and Cdc5 colocalization, the triple-mutant exhibited severe SPB fragmentation and aberrant localization of Cdc5 (**Figure 5A**). These defects extended to at least 50% of triple mutant cells and encompassed diffuse Cdc5 cellular signal, mislocalization of Cdc5 in puncta independent from SPBs, and SPB fragmentation with or without colocalization of Cdc5 (**Figure 5B**). Impaired Cdc5 colocalization did not correlate temporally with the DNA damage response, as the aberrant phenotypes were observed both before and after DNA damage induction. Importantly, the adaptation and SPB colocalization defects of the *nud1-92 spc110-7 spc72-8* mutant recapitulated the behavior of the *cdc5-16* PBD binding mutant (Ratsima et al., 2016). The fact that mislocalization of Cdc5 depends on the loss of multiple SPB components dovetails nicely with the observation that Cdc5 enriches at two spatially distinct sites on SPBs; namely, the inner (Spc110) and outer (Nud1/Spc72) plaques of the organelle (Botchkarev et al., 2017). Together, these results indicate that PBD-mediated enrichment of Cdc5 at the SPB is a precondition to maintain SPB function during adaptation to DNA damage.

**Figure 5.**
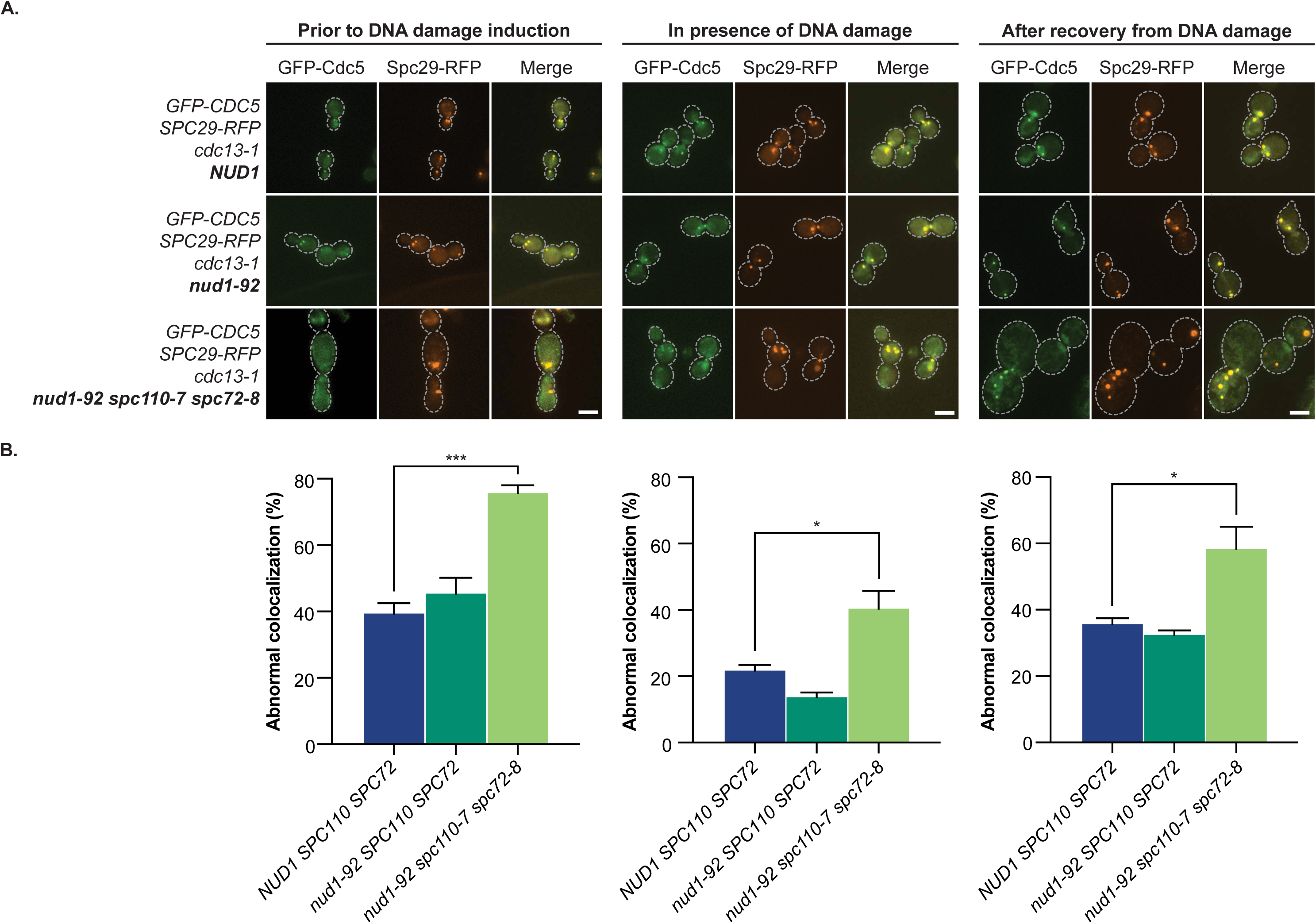
Cdc5 colocalization at the SPB depends on Nud1, Spc72 and Spc110 PBD binding sites. **(A)** A time-course experiment was performed with yeast cells carrying *cdc13-1*, *GFP-CDC5* and *SPC29-RFP* to assess Cdc5 localization at SPBs. Control cells as well as a *nud1-92* single mutant and a *nud1-92 spc110-7 spc72-8* triple-mutant were analyzed prior to DNA damage induction (left panel, room temperature), during (middle panel) and after recovery from DNA damage (right panel, room temperature). Mitotic cells were monitored for SPB morphology and Cdc5 colocalization with Spc29-RFP. Aberrant colocalization of Cdc5 at SPBs and SPB fragmentation was observed in the triple-mutant. Genotypes corresponding to each strain are shown on the left side. Scale bars correspond to 5 µm. Outlines of single cells marked with a dashed line. **(B)** Quantification of the fraction of mitotic cells showing abnormal localization of Cdc5 for the experiment shown in panel A. Phenotypes categorized as abnormal included: diffuse Cdc5 cellular signal, mislocalization of Cdc5 in GFP puncta unrelated to SPBs, and SPB fragmentation with or without colocalization of Cdc5. A minimum of 100 cells per strain was counted for each timepoint. Data represented as mean values and SEM of at least 3 independent experiments. Statistical analysis performed using a one-way ANOVA test on normally distributed values with Dunnett’s multiple comparisons post hoc test. Statistical significance is shown on graph with asterisks (* is p ≤ 0.05; *** is p ≤ 0.001).

### SPB components Cnm67 and Mps3 are substrates of Cdc5 *in vivo*

We next asked if SPB proteins are substrates of Cdc5 kinase during the adaptation response to DNA damage. To achieve this goal, we mined phosphorylation datasets (Stark et al., 2010) and analyzed SPB proteins for the presence of consensus motifs for Cdc5 kinase activity (Brar et al., 2006). Results from this analysis revealed the outer plaque component Cnm67 and the nuclear pore complex (NPC) protein Mps3 as putative substrates for Cdc5. Both Cnm67 and Mps3 share a key function in SPB dynamics by regulating SPB assembly (Brachat et al., 1998) and SPB duplication events (Jaspersen et al., 2002), respectively. To assess whether Cdc5 phosphorylates Cnm67 *in vivo*, we monitored the electrophoretic behavior of HA-tagged Cnm67 (Cnm67-3xHA) during a synchronous round of cell division in the presence and absence of functional Cdc5 kinase activity. We used a ts allele of Cdc5 defective for substrate phosphorylation (*cdc5-99;* St-Pierre et al., 2009) as well as a *cdc15-2* control (Surana et al., 1993) to allow effective comparison of the phosphorylation levels observed in each strain (i.e., both mutants have a terminal arrest in telophase). As seen in **Figure 6A**, the hyperphosphorylation shift normally detected on Cnm67-3xHA in late mitosis (120 min; left panel) was absent in the kinase-defective *cdc5-99* strain (right panel), leaving only Cdc28-mediated priming phosphorylation events on Cnm67-3xHA (Grava et al., 2006). Next, we created the *cnm67-16A* mutant, a phospho-defective allele that has lost all Cdc5 consensus sites for phosphorylation (Brar et al., 2006)(**Figure 6B**). Time-course analyses of Cnm67-3xHA and Cnm67-16A-3xHA electrophoretic behavior revealed that the Cnm67-16A mutant is completely defective for phosphorylation-induced gel retardation relative to the wild-type protein (**Figure 6C**).

**Figure 6.**
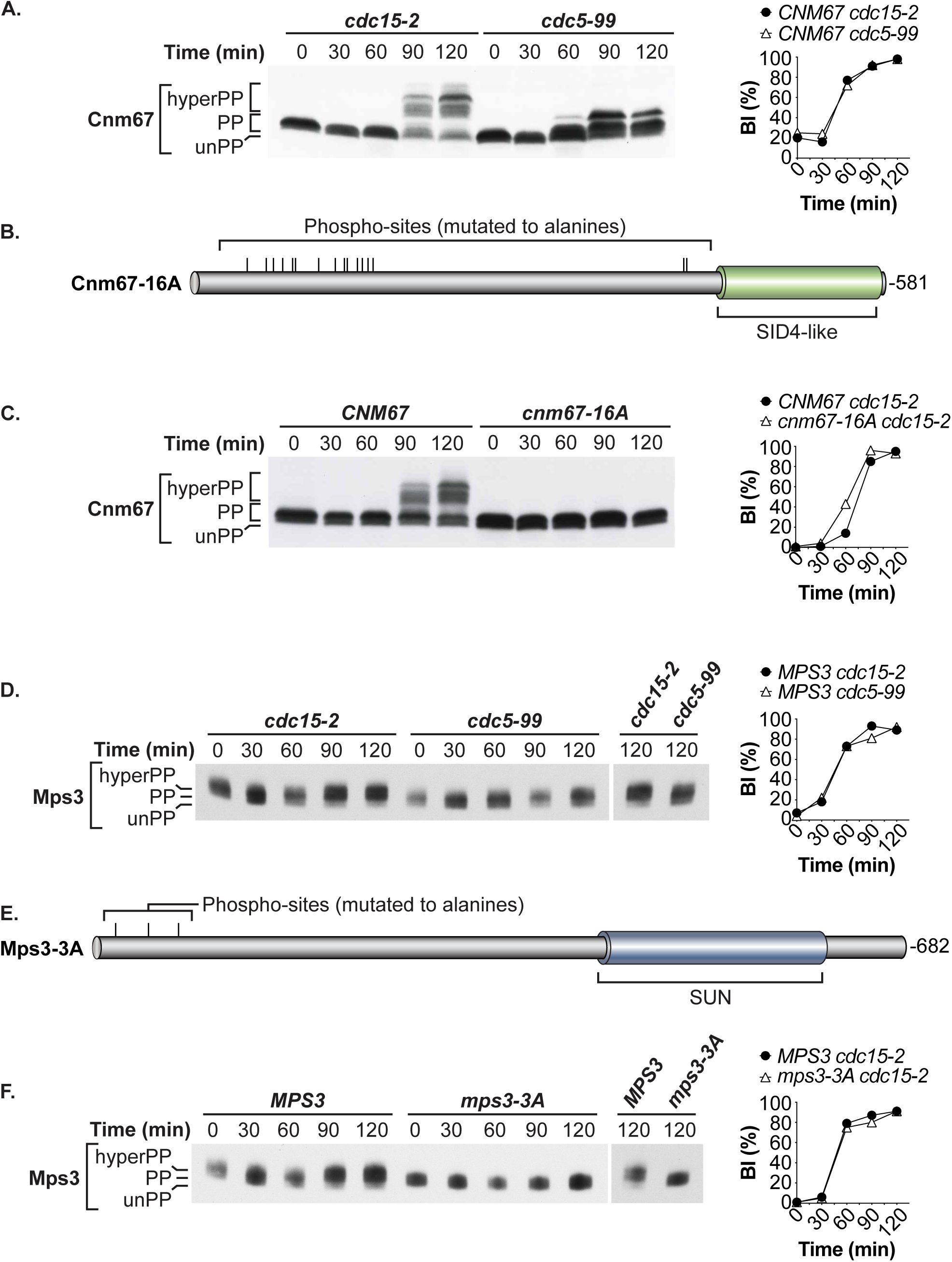
SPB components Cnm67 and Mps3 are substrates for Cdc5 *in vivo*. **(A)** Impact of Cdc5 kinase on Cnm67 phosphoryla8on. Cells synchronized in G1 with α-factor were released into a synchronous cell cycle at restric8ve temperature for *cdc5-99* and *cdc15-2* (37 °C) and samples were collected at regular intervals to monitor the electrophore8c behavior of HA-tagged Cnm67. The control yeast strain carries the *cdc15-2* muta8on to elicit a late telophase arrest similar to that of *cdc5-99* mutants (St-Pierre et al., 2009). Loss of Cnm67 hyperphosphoryla8on, as evidenced by the faster migra8on of the protein aRer SDS-PAGE, was observed in absence of func8onal Cdc5 kinase ac8vity. Budding index was monitored to assess the synchrony of cell cycle progression in cultures of *cdc15-2* and *cdc5-99* mutants (i.e., graph on the right of the immunoblots). A total of 100 cells per strain was counted for each 8me point, and a minimum of 3 independent experiments were performed. **(B)** Schema8c representa8on of phosphoryla8on site muta8ons inserted in Cnm67-16A protein. Key structural domains and posi8on of muta8ons (ver8cal lines) are represented to scale. **(C)** Cnm67 and *cnm67-16A* phosphoryla8on dynamics were compared in *cdc15-2* strains progressing synchronously in the cell cycle. Assessment of phosphoryla8on and budding index were carried out as described in panel A. All detectable forms of phosphoryla8on were lost in *cnm67-16A*-3xHA. **(D)** Mps3 phosphoryla8on status in the presence and absence of func8onal Cdc5 kinase. The experiment was performed as in panel A. **(E)** Schema8c representa8on of phosphoryla8on site muta8ons inserted in Mps3-3A protein. Other details are as described in panel B. **(F)** Phosphoryla8on state of Mps3-3A during a synchronous cell cycle. Analysis of Mps3 phosphoryla8on and cell culture synchrony are essen8ally iden8cal to the experiment described in panel A. Hyperphosphoryla8on was lost in Mps3-3A-3xHA, resul8ng in the protein being evenly distributed between an unphosphorylated state (i.e., unPP), and a basal/Cdc28-mediated phosphorylated state (i.e., PP).

Similar results were obtained when performing an analysis of phosphorylation kinetics with HA-tagged Mps3 (Mps3-3xHA). Specifically, a loss of hyperphosphorylation was detected on Mps3-3xHA in late anaphase when Cdc5’s KD activity was impaired (*cdc5-99)* (120 min; right panel - **Figure 6D**). Only basal/Cdc28-mediated phosphorylation remained on Mps3 after inactivation of Cdc5 (see “PP” isoform in Figure 6). Assessing the electrophoretic behavior of the Mps3-3A phospho-mutant (**Figure 6E**) after SDS-PAGE revealed its inability to undergo hyperphosphorylation, as all detectable Mps3-3A-3xHA species were evenly distributed between an unphosphorylated state (i.e., unPP) and a basal/Cdc28-mediated phosphorylated state (i.e., PP; **Figure 6F**; Prasada Rao et al., 2021; Ubersax et al., 2003). Taken together, these results show that effective phosphorylation of Cnm67 and Mps3 requires full Cdc5 activity.

### Phosphorylation of Cnm67 and Mps3 is required to induce an adaptation response to DNA damage

The observation that Cdc5 modulates Cnm67 and Mps3 phosphorylation levels at SPBs suggests that these modifications might be physiologically relevant for the adaptation response to DNA damage. To test this notion, we performed a microcolony formation assay with *cnm67-16A* and *mps3-3A* mutants carrying *cdc13-1* (**Figure 7B**). Both *cnm67-16A* and *mps3-3A* were impaired for microcolony formation in response to DNA damage induction, with a 30%–40% reduction relative to control after 24 hours of growth (**Figure 7C**). The adaptation response to persistent DNA damage is therefore compromised in *cnm67-16A* and *mps3-3A* phospho-mutants. Similar results were obtained when testing microcolony formation in response to a sustained HO-induced DSB (**Figure S4**). Importantly, the *cnm67-16A* and *mps3-3A* phospho-defective alleles do not show detectable proliferation defects under a wide range of temperatures (**Figure 7A**), indicating that they are proficient in the essential MTOC function of SPBs and their adaptation defect is therefore mechanistically distinct from the core function of SPB in microtubule nucleation.

**Figure 7.**
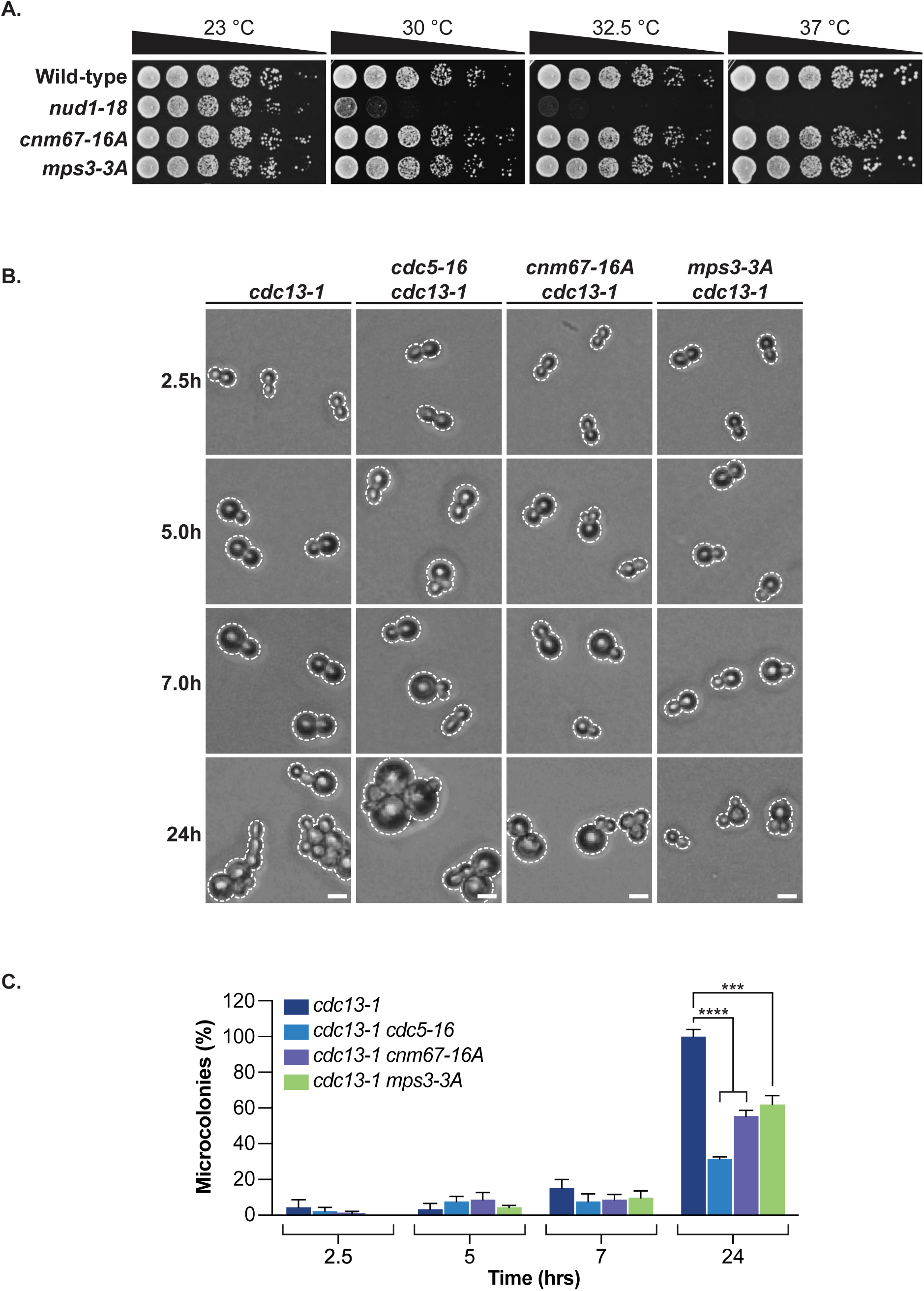
Cdc5-dependent phosphorylation of Cnm67 and Mps3 are required to signal adaptation to DNA damage. **(A)** Proliferation of phospho-mutants at various temperatures was assessed by serial dilution assay on solid media and growth at 23 °C, 30 °C, 32.5 °C and 37 °C. *cnm67-16A* and *mps3-3A* phospho-mutants proliferated normally at all temperatures tested. (n=3) **(B)** The ability of *cnm67-16A* and *mps3-3A* phospho-mutants to elicit an adaptation response after inducing *cdc13*-dependent/unrepairable DNA damage was assessed using a standard microcolony formation assay. c*nm67-16A* and *mps3-3A* both showed defects in adaptation to DNA damage relative to control strain. Scale bars correspond to 10 µm. Outlines of cells and microcolonies marked with a dashed line. **(C)** Quantification of the fraction of individual cells growing into microcolonies for the strains tested in panel B. A minimum of 100 cells per strain was counted for each timepoint. All values were normalized to 100 based on the T24 h mean value of the *cdc13-1* control strain. Data represented as normalized mean values and SEM of at least 3 independent experiments. Statistical analysis performed using a one-way ANOVA test on normally distributed values with Dunnett’s multiple comparisons post hoc test. Statistical significance is shown on graph with asterisks (*** is p ≤ 0.001; **** is p ≤ 0.0001).

## Discussion

The capacity of cells to proliferate in the presence of unrepairable DNA damage is an intriguing biological property from the perspective of genome stability and organism homeostasis. While the adaptation pathway promotes cell survival in the short term, it also increases the risk of genome instability, which is an underlying cause for cellular dysfunction and cancer development. In this context, it appears likely that the adaptation response to DNA damage has been finely honed by evolution to optimize cell survival without dramatically increasing cancer occurrence in multicellular organisms. Polo kinases are the most important regulators of this adaptation response in eukaryotes, and their activity is tightly regulated to ensure the careful balance between cell survival and organism homeostasis. We have recently shown that centrosomes are key components of yeast PLK/Cdc5 kinase regulation during the cellular response to DNA damage (Ratsima et al., 2016). Until now, the issue of how SPBs contribute to cellular adaptation to DNA damage has remained a key unanswered question. We found in this study that Cdc5 takes advantage of essential SPB components to promote DNA damage adaptation, adding unexpected players and mechanistic insights to our understanding of Cdc5’s role in this pathway. In particular, our findings shed light on the nature of Cdc5 interactome at SPBs and highlight how essential processes involved in cell cycle progression are modulated to promote cell survival under challenging conditions for genome stability.

Our study revealed a central role for Nud1 as a platform that coordinates the action of Cdc5 at SPBs during the adaptation response to DNA damage. What might be the extent of Nud1 input in this process? Nud1 may contribute to this pathway by 1) recruiting Cdc5 at SPBs to facilitate transduction of a crucial signal that overrides DNA damage checkpoint activation, and 2) acting as a MEN scaffold that induces mitotic exit when the G2/M checkpoint is silenced and cells become receptive to cell cycle re-entry with lingering DNA damage (Hu et al., 2001; Valerio-Santiago et al., 2013). The observation that a hyperactive Nud1 allele (i.e., *nud1-A308T)* can trigger partial spindle assembly checkpoint (SAC) bypass and robust SPoC bypass through Cdc5-independent constitutive recruitment of the Hippo-like Cdc15 kinase is consistent with this hypothesis (Vannini et al., 2021). In addition, it is possible that other SPB receptors for Cdc5 –like Spc72– would contribute to the adaptation response via their roles in the sequential and asymmetric recruitment of SPB components to the organelle (Matellán et al., 2020).

Results obtained from our genetic interaction analyses with the SPB triple-mutant *nud1-92 spc110-7 spc72-8* confirmed that Cdc5 uses more than one recruitment factor to mediate its enrichment at the SPB during DNA damage adaptation. Mutating PBD-binding sites in the triple-mutant significantly exacerbated *nud1-92* adaptation phenotype and led to microcolony formation defects equivalent in severity to the ones of the PBD-defective allele *cdc5-16*, without impairing essential cell division processes. It is noteworthy that the essential functions of SPB in MEN and MTOC pathways were maintained in *nud1, spc110, and spc72* PBD mutants, likely reflecting a lower concentration/activity threshold required for these proteins to maintain MEN activation and spindle formation relative to the levels necessary to promote DNA damage adaptation. Assessing Cdc5-SPB colocalization in the PBD triple-mutant demonstrated that Cdc5 requires a significant number of independent PBD-binding motifs to mediate its effective recruitment to SPBs, implying a multivalent mode of interaction based on weak and/or transient associations with several PBD-binding sites at the organelle. Consistent with this view, Cdc5 colocalization with SPBs was only impaired when a substantial number of PBD-binding sites were removed from Nud1, Spc110 and Spc72. Importantly, this colocalization defect was associated with aberrant SPB foci formation (with or without Cdc5 colocalization), diffuse Cdc5 cellular signal, and mislocalization of Cdc5 in aggregates distinct from SPBs. Together, these observations suggest that multi-site recruitment of Cdc5 at SPBs is important to mediate and/or modulate the function of PLKs at centrosomes.

As Nud1 behaves as a sensor for cytoplasmic microtubule organization throughout the cell cycle and actively contributes to mitotic exit via the MEN (Gruneberg et al., 2000), it is tempting to speculate that Cdc5 might fine-tune cell cycle progression by interacting with a fixed group of SPB components to coordinate spatial cues with the timing of mitotic events both under normal conditions and upon checkpoint activation. The identification of Spc110 and Spc72 as additional Cdc5 SPB scaffolds/interactors during DNA damage adaptation supports this notion. While Spc110 regulates mitotic spindle formation (Stirling et al., 1996) and promotes timely mitotic exit (Abbasi et al., 2022), Spc72 is a key SPB component required for spindle function (Souès and Adams, 1998) and SPoC regulation (Maekawa et al., 2007). Cdc5 also phosphorylates the SAC factor Mad3 at kinetochores under normal conditions and especially upon SAC activation (Rancati et al., 2005), suggesting that the pattern of Cdc5 spatiotemporal enrichment across the cell might remain the same regardless of checkpoint stimulus. Such a biological system would facilitate Cdc5 dynamic activity in promoting mitotic progression regardless of cycling conditions, or the nature of the checkpoint involved.

Our work demonstrates that SPBs actively partake in the regulation of cell division under strenuous DNA damage conditions, which falls directly in line with the essential role of these organelles as signal transduction organizing centers (STOCs; Arquint et al., 2014). Previous work has established centrosomes/SPBs as central hubs capable of coordinating complex signaling events via transient recruitment of kinases, cell cycle regulators, and cell fate effectors relevant to diverse biological functions (reviewed in Langlois-Lemay and D’Amours, 2022). In the context of our work, it is remarkable that eukaryotic cells have leveraged SPBs/centrosomes as STOCs to coordinate the complex sequence of events necessary to bypass checkpoint arrest. Cdc5-mediated phosphorylation of the canonical MEN substrate Bfa1 represents another good example of the STOC role of SPBs relevant to DNA damage conditions (Hu et al., 2001; Valerio-Santiago et al., 2013). Taken together with previous work (Ratsima et al., 2016), our study establishes centrosomes/SPBs as active participants in cell fate decisions leading to cell cycle re-entry in the presence of persistent DNA damage. Our data supports a model whereby SPBs act as structural platforms on which spatial and temporal cues relevant to DNA damage are integrated and signalled to downstream cell cycle effectors. Overall, our findings shed light on the nature of Cdc5 interactome at SPBs and highlight how essential processes involved in cell cycle progression are modulated to promote cell survival under DNA damage conditions. Gaining a better understanding of how PLK function is regulated at centrosomes and its impact on the proliferation of cells carrying unrepairable DNA damage has important ramifications for human health. For instance, the poor prognosis associated with PLK1-overexpressing cancers (reviewed in Liu et al., 2017; Chiappa et al., 2022; Gheghiani and Fu, 2023) and their acute resistance to radiotherapy (Syljuåsen et al., 2006; Rödel et al., 2010; Hagege et al., 2021) or drug treatments (Sero et al., 2014) may be directly linked to the enhanced recruitment of PLK1 observed at centrosomes in cancer cells and tumor tissues (Xie et al., 2020). It is tempting to speculate that some of the tumorigenic effects associated with *PLK1* overexpression is the consequence of an enhanced adaptation response to DNA damage, similar to the phenotype observed with *CDC5*-overexpression in yeast cells. In this context, the development of inhibitors specifically designed to impede PLK1 action at centrosomes might be effective in preventing tumor cell proliferation in the presence of radiotherapy/drug-induced DNA damage. Future work in mammalian cell models will determine if this therapeutic strategy holds promise for the treatment of human cancers.

## Materials and Methods

All yeast strains used in this study are derived from a W303 background (see Table S1 for a detailed list of strain genotypes). For yeast culture and genetic manipulations (including mating, sporulation and dissection), standard procedures were used (Guthrie and Fink, 1991). For the generation of mutant alleles as well as epitope tagging, PCR amplification and yeast transformation were used as detailed in previous literature (Longtine et al., 1998). Standard conditions were used to conduct microcolony formation assays (Toczyski et al., 1997; Lee et al., 1998b; Vidanes et al., 2010), as well as electrophoresis and immuno-blotting procedures (Foiani et al., 1994; D’Amours and Jackson, 2001; St-Pierre et al., 2009). All quantifications in this study represent the means ± SEM of at least three independent replicates. GraphPad Prism 9 for MacOS (version 9.5.1) was used to process data and run all statistical tests. An extensive list of all materials and methods can be found in Supplemental experimental procedures.

## Competing Interest Statement

The authors declare no competing interests.

## Acknowledgements

We would like to thank Dr. Malika Saint for helpful comments on the manuscript. Research in D.D.’s laboratory is supported by a grant from CIHR (FDN–167265) and by the Canada Research Chair in Chromatin Dynamics & Genome Architecture (CRC-2017-00064).

## Author Contributions

L.L.L. and D.D. conceived and designed the experiments. L.L.L. performed the experiments; L.L.L. and D.D. analyzed the Data; L.L.L. and D.D. wrote the paper.

## Supplemental Information

Supplemental information includes one table detailing yeast strains used in this study, Supplemental experimental procedures, and four Supplementary figures (S1-S4).

## Supplemental Experimental Procedures

### Yeast proliferation assays

Strains were incubated overnight at room temperature (23 °C), diluted to 0.25 OD_600nm_ the next morning and grown until they reached exponential phase at permissive temperature (23 °C). Five-fold dilution series with the first dilution corresponding to an OD_600nm_ of 0.2 were spotted on solid YEPD medium (yeast extract, peptone, D-glucose) and grown in temperature-controlled incubators (Binder) for 24-96 hours prior to scanning and processing.

### Microcolony formation assays

To monitor yeast cell adaptation to unrepairable DNA damage, we performed microcolony assays under DNA damage conditions induced by inactivation of the telomere-capping protein Cdc13 (*i.e., cdc13-1*) at restrictive temperature (between 31 °C and 33 °C; Weinert et al., 1994). Cells were inoculated in liquid YEPD medium, grown at room temperature overnight and diluted to 0.25 OD_600nm_ the next morning. After reaching mid-log phase (0.3 OD_600nm_), cultures were transferred into a shaking water bath at restrictive temperature (T=0h) for a duration of 2 hours to activate the DNA damage checkpoint in response to *cdc13-1* inactivation. Then, each culture was plated on the surface of 4 YEPD plates (one for each time point analyzed) in a temperature-controlled room to maintain DNA damage-inducing conditions. Stacks for each tested strain comprised 4 plates (plates: T=2.5h, T=5h, T=7h, T=24h) and were incubated at restrictive temperature for the indicated time frame. Cell populations in each plate were assessed for DNA damage adaptation at its corresponding time point. Cell morphology was documented using digital microscopic photography on a tetrad dissection microscope (Nikon *Eclipse 50i*) equipped with a DS-Fi1 camera (Nikon). Raw images were adjusted using the Fiji/ImageJ software (National Institutes of Health) to optimize brightness and contrast. Microcolonies were defined as an interconnected group of at least 5 cell bodies with less than half of the cells displaying a diameter above 10 μm. A minimum of 100 cells per strain was counted in each replicate. In all cases, a minimum of three independent experiments were performed.

In microcolony assays in which *CDC5* was overexpressed, the same experimental procedure was used with a minor modification. Cells were incubated overnight in raffinose-containing medium (YEPR – 2%) and diluted in YEPR the next morning. Cells were plated on solid YEP medium containing 2% galactose (YEPG).

In microcolony assays using the HO endonuclease, cells were genetically modified to generate a sustained double-strand break using an inducible *GAL1* promoter to modulate HO expression and activity (*P_GAL1_-HO*; as described in Lee et al., 1998b). In these assays, cultures of cells were grown overnight at room temperature and diluted the next morning at 0.25 OD_600nm_ in fresh YEP-2% Raffinose medium. Cells were grown until they reached mid-log phase, after which each culture was directly plated on the surface of solid YEP-2% Galactose medium (T=0h) and monitored for microcolony formation at various timepoints (T=2h, T=6h, T=24h, T=48h). The different duration and sample collection timeframes of this adaptation experiment compared to the *cdc13-1* protocol reflects the differences in proliferation kinetics of yeast cells in the presence of galactose and HO-induced DNA damage. Throughout the HO adaptation experiments, cells were maintained at a temperature of 23 °C. For *cnm67-16A*, cells were placed with a dissection needle in a grid-like arrangement on the surface of solid medium and monitored for adaptation rates (Lee et al., 1998b). The same plate was monitored at each timepoint to assess microcolony formation. Prior to adaptation tests, “micro-manipulation” survival ratios were established for the control strain and *cnm67-16A* mutant by plating 100 cells of each strain with a dissection needle on solid YEPD media (i.e., without DNA damage) and assessing microcolony formation after 6 hours of growth. This control was necessary to determine if yeast strains were sensitive to needle manipulation. Survival to needle manipulation was defined as the mean value of the percentage of cells surviving needle plating from three independent experiments. Values obtained in the adaptation tests were normalized to the micro-manipulation survival ratio to accurately reflect rates of adaptation. Survival rates in response to the physical stress induced by the dissection needle in *nud1* and *mps3-3A* mutants were too low to yield quantifiable survival values. To circumvent this limitation, *nud1* and *mps3-3A* mutants were plated onto solid medium using a traditional carpet-plating technique and analyzed for adaptation to persistent DNA damage as previously described.

When applicable, values were normalized to 100% based on the T24 h or T48 h mean value of the control strain/strain showing the highest mean value to maintain comparable results across experiments and facilitate visual comparisons between figures. Figure legends are reflective of data normalization when applied.

### Protein phosphorylation analysis

The phosphorylation levels of Cnm67-3xHA, Cnm67-16A-3xHA, Mps3-3xHA and Mps3-3A-3xHA were assessed after electrophoresis and immuno-blotting (St-Pierre et al., 2009). Total cell extracts for each timepoint were processed using the TCA glass beads method (Foiani et al., 1994) and loaded on a 10% homemade low-bis 150:1 ratio gel (D’Amours and Jackson, 2001). Proteins were transferred onto a PVDF membrane using a wet transfer procedure and visualized by immuno-blotting using the mouse monoclonal antibody 12CA5 (Cnm67-3xHA and Cnm67-16A-3xHA) (Roche, #11583816001) at a dilution of 1:5000 in PBS-T supplemented with 3% milk, or the 16B12 epitope (Mps3-3xHA and Mps3-3A-3xHA) at a dilution of 1:2500 in PBS-T supplemented with 3% milk. Small samples of each cell culture were collected (in parallel to samples dedicated to protein analysis) to determine the budding index (BI) as cells progressed in the time-course experiment. Specifically, fractions of the cell cultures (10 ml per timepoint) were pelleted and resuspended in 250 μl 20% TCA. A volume of 50 μl was aliquoted into a separate tube, pelleted and resuspended in 1 ml 70% ethanol (dedicated to BI counting) while the remaining 200 μl was processed for electrophoresis and immuno-blotting using the TCA glass beads method. The fraction of budded cells at each time-point was counted on a light microscope (Nikon *Eclipse 50i*), as previously described (St-Pierre et al., 2009).

### Fluorescence microscopy

To monitor SPB dynamics and Cdc5 colocalization *in vivo*, Cdc5-GFP and Spc29-RFP markers were introduced in *nud1-92* and *nud1-92 spc110-7 spc72-8* mutant strains carrying *cdc13-1*. The colocalization of Cdc5-GFP and Spc29-RFP was assessed in a time-course experiment prior to, during, and after recovery from DNA damage induction. Briefly, cells were grown overnight and diluted at 0.25 OD_600nm_ the next morning at room temperature (23 °C). After two hours at 23 °C, the first sample (T=0h – before DNA damage induction; 1 ml) was aliquoted and cultures were transferred into a shaking water bath for 2 hours at restrictive temperature to inactivate *cdc13-1* and generate DNA damage. After 2 hours in the water bath, a second sample (T=2h – during DNA damage; 1 ml) was collected. Then, cultures were transferred at room temperature and grown for another 2 hours before collecting the final sample (T=4h – after recovery from DNA damage; 1 ml). Cell culture samples were processed for fluorescence microscopy imaging using a standard procedure. Briefly, cells were pelleted and resuspended in a 0.1 M potassium phosphate solution (pH 6.4) containing 3.7% of formaldehyde. Cells were fixed in this solution for 20 min. Samples were then washed twice with a 0.1 M potassium phosphate solution (pH 7). Final samples were resuspended in 100 μl potassium phosphate buffer (pH 7) and 5 μl was spotted on a microscope slide for observation. Samples were analyzed using a Nikon Eclipse T*i*2 inverted microscope equipped with a 100X objective. A minimum of 100 cells were scored for each timepoint and SPB behavior/Cdc5-SPB colocalization was monitored using *NIS elements* software (Nikon Instruments Inc., NY, USA). Phenotypes categorized as abnormal included: Diffuse Cdc5 cellular signal, mislocalization of Cdc5 in GFP puncta unrelated to SPBs, and SPB fragmentation with or without colocalization of Cdc5. Raw images were processed, and scale bars were added using Fiji/ImageJ.

### Statistical analyses

All statistical analyses were performed using GraphPad Prism 9 for macOS (version 9.5.1). Normal data distribution was assessed using the Shapiro-Wilk test in which a W ≥ 0.7500 represented data fitting standard normal quantiles. Q-Q plots were analyzed and used as an auxiliary supporting method to evaluate data distribution. To test for statistical significance, an ordinary one-way ANOVA was performed on normally distributed values followed by a Dunnett’s multiple comparisons post hoc test to report statistically significant differences between control group and mutants. When only two groups were compared, an unpaired t-test on normally distributed values was performed. Statistical significance was represented as asterisks on graphs (* for p ≤ 0.05; ** for p ≤ 0.01; *** for p ≤ 0.001; **** for p ≤ 0.0001).

---

**Figure S1.**
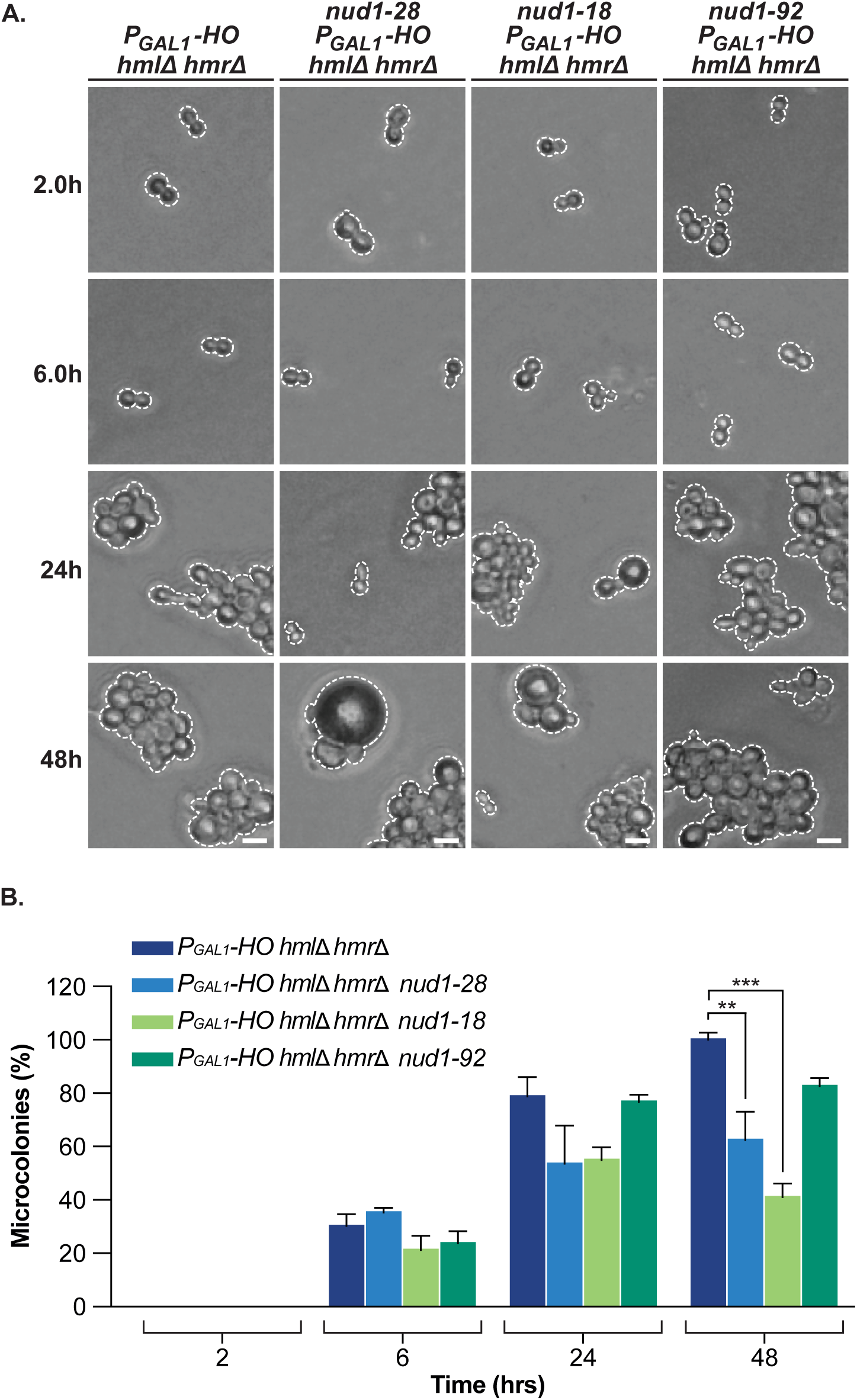
*nud1* mutants are defective for adaptation to a sustained HO-induced double-strand break (DSB). **(A)** Microcolony formation assay performed at 23 °C using overexpression of HO endonuclease (*P_GAL1_-HO*) as a source of persistent DNA damage. Strains were grown in YEPR (2% raffinose) and plated on solid YEPG (2% galactose) to induce DNA damage (see methods for a detailed description of the assay). Rates of microcolony formation following DNA damage induction were lower in *nud1* mutants relative to control strain. Scale bars correspond to 10 µm. Outlines of cells and microcolonies marked with a dashed line. Differences in genotypes for each strain described in panel headings. **(B)** Quantification of the fraction of cells growing into microcolonies for each strain and timepoint shown in panel A. A minimum of 100 cells per strain was counted for each timepoint. All values were normalized to 100 based on the T48 h mean value of the control strain. Data represented as normalized mean values and SEM of at least 3 independent experiments. Statistical analysis performed using a one-way ANOVA test on normally distributed values with Dunnett’s multiple comparisons post hoc test. Statistical significance is shown on graph with asterisks (** is p ≤ 0.01; *** is p ≤ 0.001).

**Figure S2.**
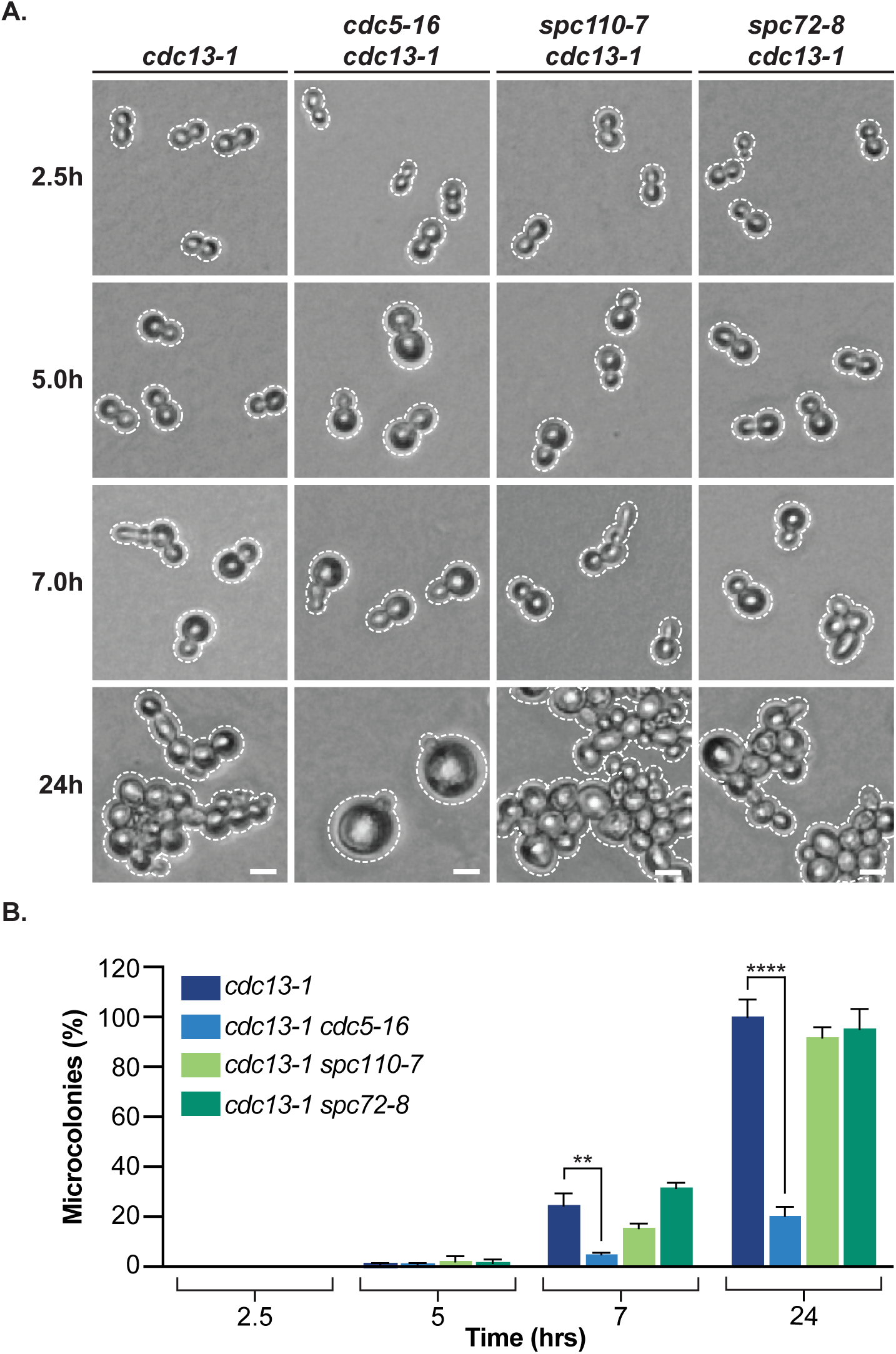
Strains carrying *spc110-7* or *spc72-8* alleles are functional for adaptation to DNA damage. **(A)** Mutant strains carrying *spc110-7* or *spc72-8* were assayed for microcolony formation following DNA damage induction at 31 °C. Both mutants formed microcolonies with kinetics mirroring a *cdc13-1* control. Scale bars, genotype notations and cell outlines are as described in Figure S1. **(B)** Quantification of the fraction of cells growing into microcolonies for each timepoint in strains tested in panel A. Data represented as normalized mean values and SEM of at least 3 independent experiments. Statistical analysis performed using a one-way ANOVA test on normally distributed values with Dunnett’s multiple comparisons post hoc test. Statistical significance is shown on graph with asterisks (** is p ≤ 0.01; **** is p ≤ 0.0001).

**Figure S3.**
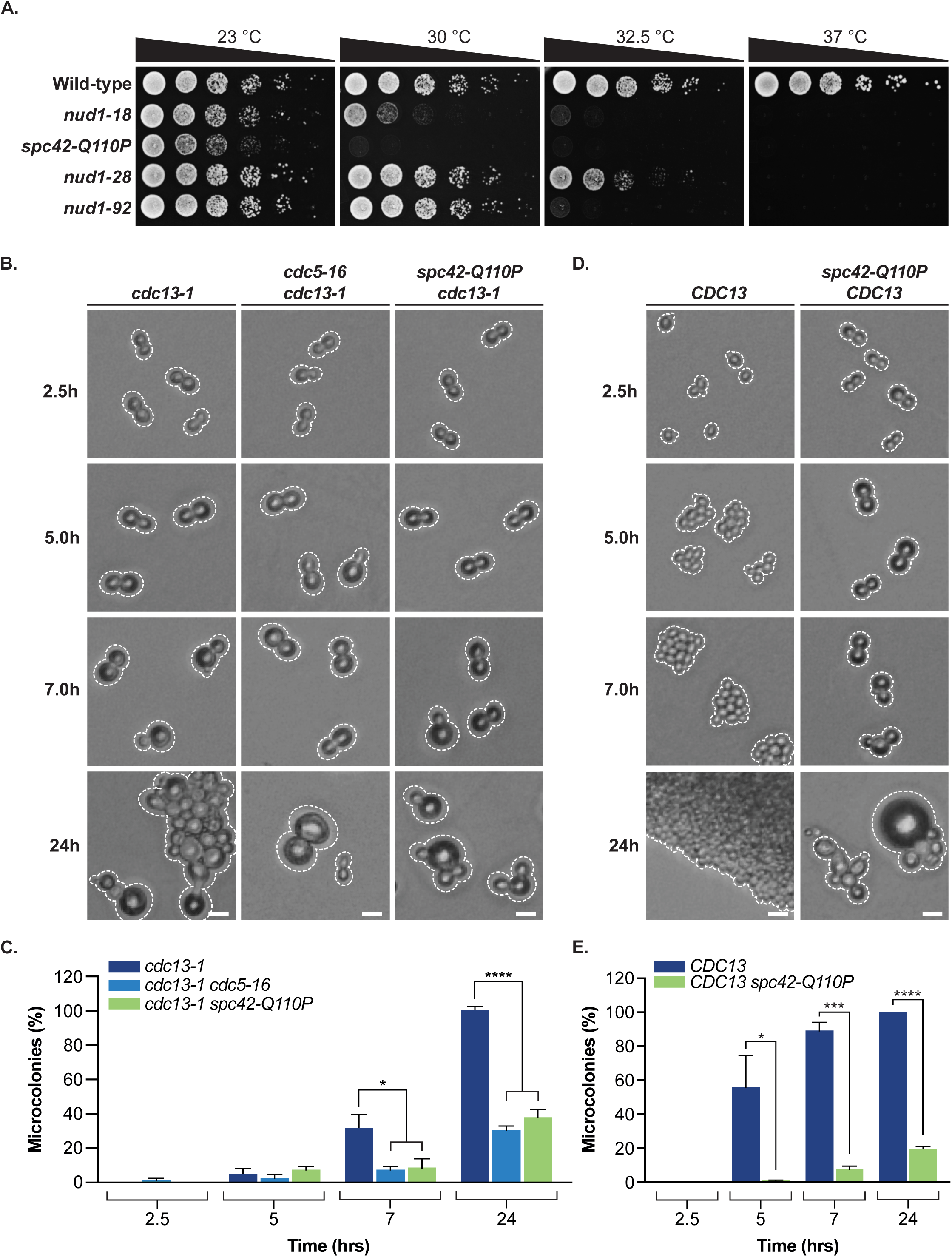
Generic inactivation of MTOC activity is not associated with a specific defect in adaptation to DNA damage. **(A)** Growth properties of a yeast mutant carrying *spc42-Q110P*. Cell cultures were serially diluted, spotted on solid medium and grown at 23 °C, 30 °C, 32.5 °C and 37 °C. The *spc42-Q110P* mutant demonstrated a severe temperature-sensitive growth phenotype. (n=3) **(B)** A yeast strain expressing *spc42-Q110P* was assayed for microcolony formation after inactivation of *cdc13-1* at restrictive temperature (32.5 °C). Scale bars, genotype notations and cell outlines are as described in Figure S1. **(C)** Quantification of the fraction of cells growing into microcolonies for each timepoint in strains tested in panel B. Data represented as normalized mean values and SEM of at least 3 independent experiments. Statistical analysis performed using a one-way ANOVA test on normally distributed values with Dunnett’s multiple comparisons post hoc test. Statistical significance is shown on graph with asterisks (* is p ≤ 0.05; **** is p ≤ 0.0001). **(D)** Microcolony formation ability of the *spc42-Q110P* mutant growing at 32.5 °C in the absence of DNA damage (*CDC13*). Notice how a global inactivation of SPB/MTOC activity causes a permanent mitotic arrest independently of DNA damage in the *spc42-Q110P* mutant. A similar defect was not observed in *nud1* mutants. **(E)** Quantification of the fraction of *CDC13* cells forming microcolonies at each timepoint of the experiment shown in panel D. A minimum of 100 cells per strain was counted for each timepoint. Data represented as mean values and SEM of at least 3 independent experiments. Statistical analysis performed using a one-way ANOVA test on normally distributed values with Dunnett’s multiple comparisons post hoc test. Statistical significance is shown on graph with asterisks (* is p ≤ 0.05; *** is p ≤ 0.001; **** is p ≤ 0.0001).

**Figure S4.**
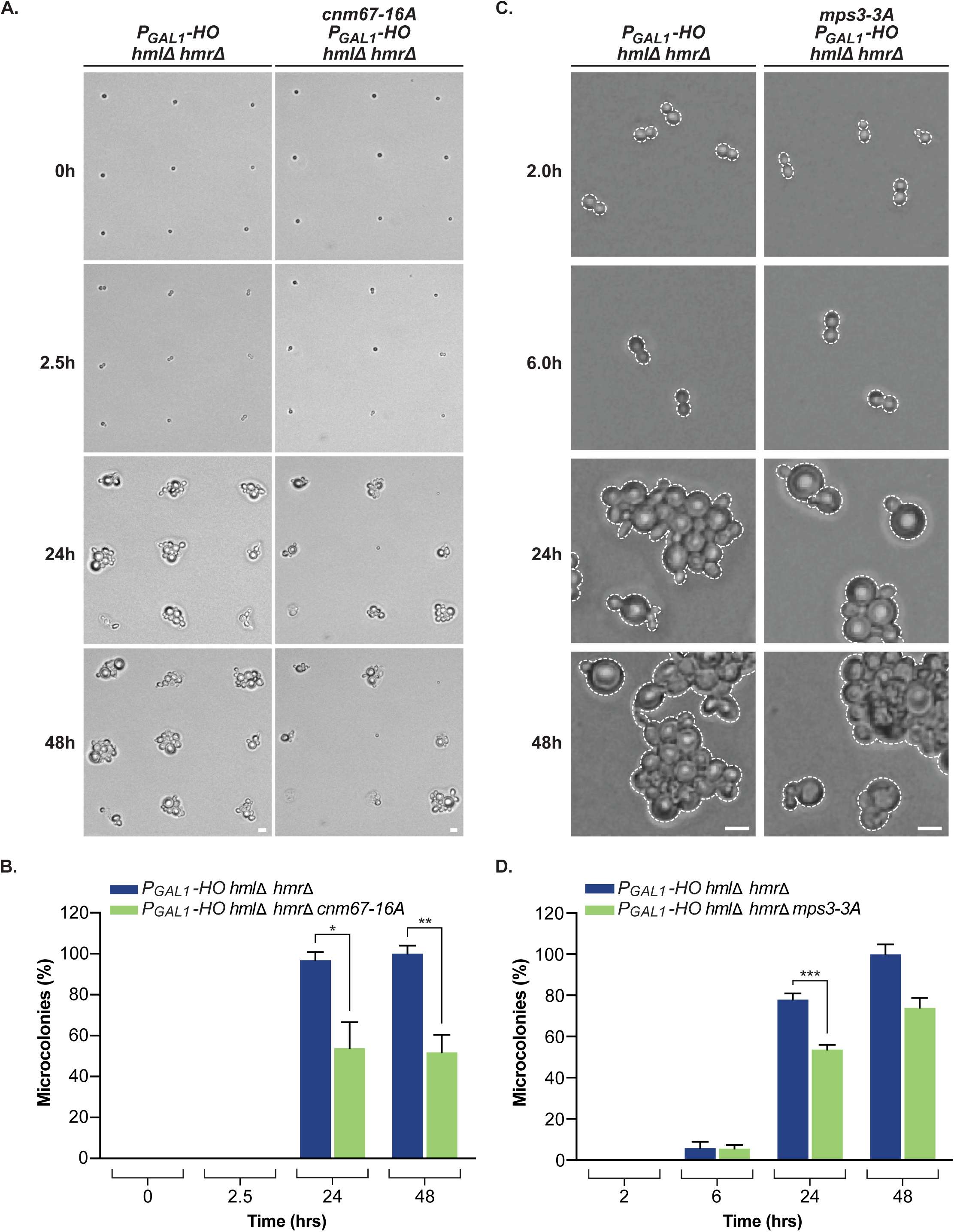
*cnm67-16A* and *mps3-3A* SPB phospho-mutants are defective for the adaptation response to a sustained double-strand break (DSB). **(A)** Microcolony formation in the presence of persistent DNA damage induced by HO endonuclease (*P_GAL1_-HO*). Strains were grown in YEPR (2% raffinose) and plated on solid YEPG (2% galactose) to induce DNA damage. 100 cells were plated in a grid-like pattern on the surface of solid medium using a dissection needle. The ability of *cnm67-16A* to adapt to a persistent DSB was reduced relative to control strain. Scale bars, genotype notations and cell outlines are as described in Figure S1. **(B)** Quantification of the fraction of individual cells growing into microcolonies for each timepoint in the strains shown in panel A. A minimum of 100 cells per strain was counted for each timepoint. All values are normalized to a relevant micro-manipulation survival ratio (see methods for details) and expressed on a percent basis relative to the value obtained at the T48 h time-point for the control strain. Data represented as normalized mean values and SEM of at least 3 independent experiments. Statistical analysis performed using a one-way ANOVA test on normally distributed values with Dunnett’s multiple comparisons post hoc test. Statistical significance is shown on graph with asterisks (* is p ≤ 0.05; ** is p ≤ 0.01). **(C)** Microcolony formation assay in the presence of HO endonuclease (*P_GAL1_-HO*) overexpression at 23° C. Strains were grown in YEPR (2% raffinose) and plated on solid YEPG (2% galactose) to induce DNA damage. Rates of microcolony formation following DNA damage induction were lower in *mps3-3A* relative to the control strain. **(D)** Quantification of microcolony formation for the strains tested in panel C. A minimum of 100 cells per strain was counted for each timepoint. All values were normalized to 100% based on the T48 h mean value of the control strain. Data represented as normalized mean values and SEM of at least 3 independent experiments. Statistical analysis performed using a one-way ANOVA test on normally distributed values with Dunnett’s multiple comparisons post hoc test. Statistical significance is shown on graph with asterisks (*** is p ≤ 0.001).

**Table S1.**
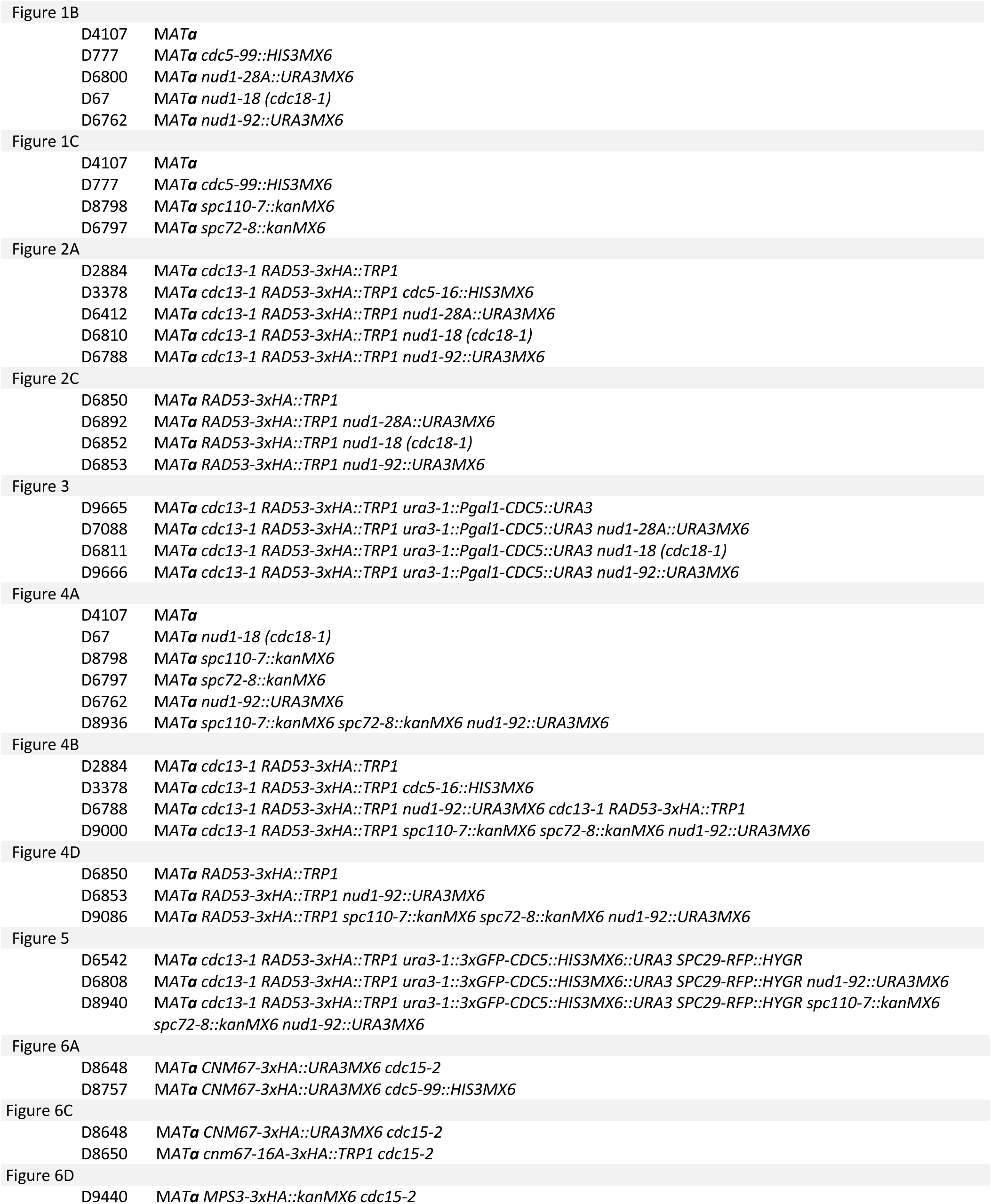
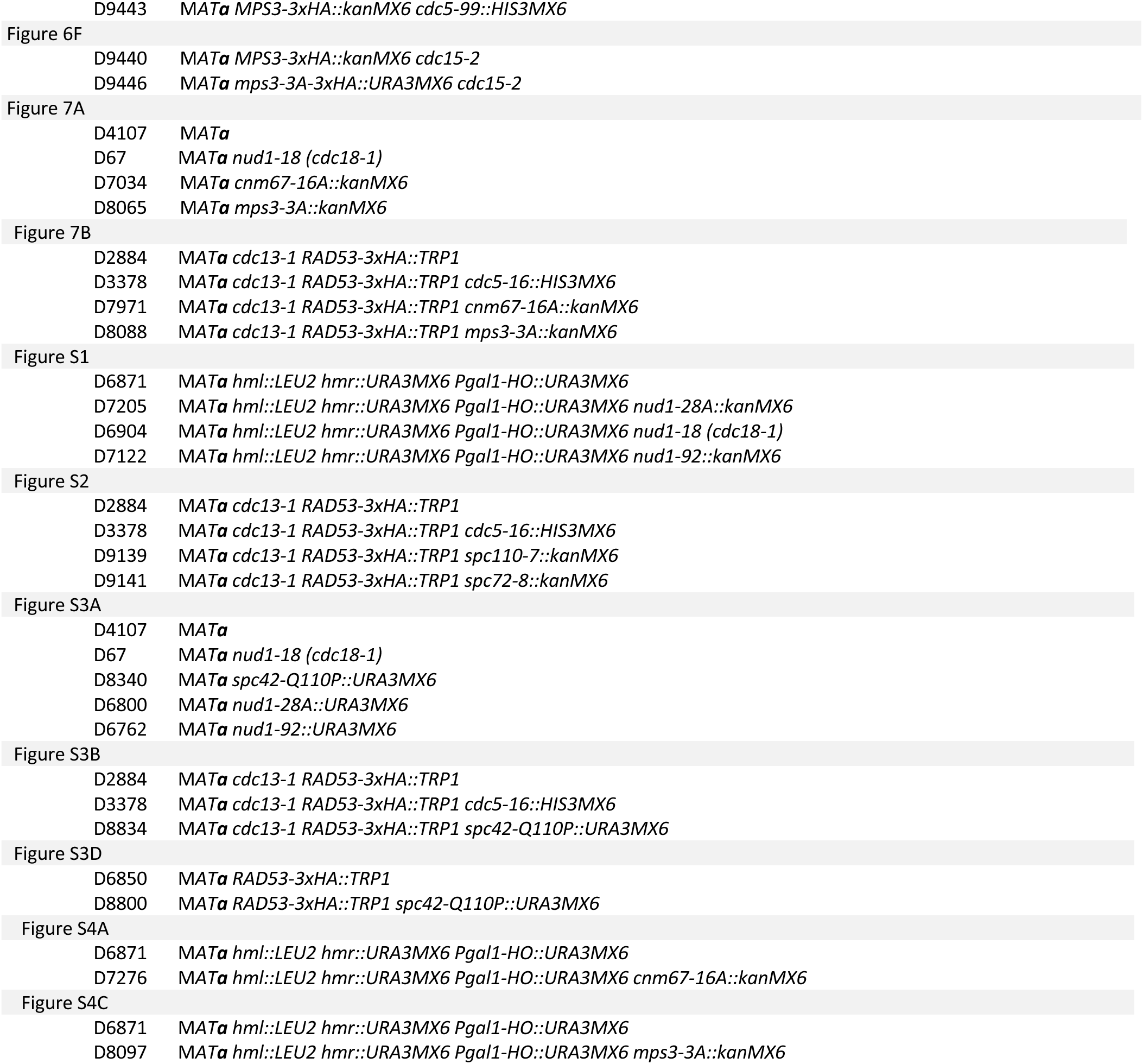
Yeast strains used in this study.

|  |  |
| --- | --- |
| Figure 1B |  |
| D4107 | MATa |
| D777 | MATa *cdc5-99::HIS3MX6* |
| D6800 | MATa *nud1-28A::URA3MX6* |
| D67 | MATa *nud1-18 (cdc18-1)* |
| D6762 | MATa *nud1-92::URA3MX6* |
| Figure 1C |  |
| D4107 | MATa |
| D777 | MATa *cdc5-99::HIS3MX6* |
| D8798 | MATa *spc110-7::kanMX6* |
| D6797 | MATa *spc72-8::kanMX6* |
| Figure 2A |  |
| D2884 | MATa *cdc13-1 RAD53-3xHA::TRP1* |
| D3378 | MATa *cdc13-1 RAD53-3xHA::TRP1 cdc5-16::HIS3MX6* |
| D6412 | MATa *cdc13-1 RAD53-3xHA::TRP1 nud1-28A::URA3MX6* |
| D6810 | MATa *cdc13-1 RAD53-3xHA::TRP1 nud1-18 (cdc18-1)* |
| D6788 | MATa *cdc13-1 RAD53-3xHA::TRP1 nud1-92::URA3MX6* |
| Figure 2C |  |
| D6850 | MATa *RAD53-3xHA::TRP1* |
| D6892 | MATa *RAD53-3xHA::TRP1 nud1-28A::URA3MX6* |
| D6852 | MATa *RAD53-3xHA::TRP1 nud1-18 (cdc18-1)* |
| D6853 | MATa *RAD53-3xHA::TRP1 nud1-92::URA3MX6* |
| Figure 3 |  |
| D9665 | MATa *cdc13-1 RAD53-3xHA::TRP1 ura3-1::Pgal1-CDC5::URA3* |
| D7088 | MATa *cdc13-1 RAD53-3xHA::TRP1 ura3-1::Pgal1-CDC5::URA3 nud1-28A::URA3MX6* |
| D6811 | MATa *cdc13-1 RAD53-3xHA::TRP1 ura3-1::Pgal1-CDC5::URA3 nud1-18 (cdc18-1)* |
| D9666 | MATa *cdc13-1 RAD53-3xHA::TRP1 ura3-1::Pgal1-CDC5::URA3 nud1-92::URA3MX6* |
| Figure 4A |  |
| D4107 | MATa |
| D67 | MATa *nud1-18 (cdc18-1)* |
| D8798 | MATa *spc110-7::kanMX6* |
| D6797 | MATa *spc72-8::kanMX6* |
| D6762 | MATa *nud1-92::URA3MX6* |
| D8936 | MATa *spc110-7::kanMX6 spc72-8::kanMX6 nud1-92::URA3MX6* |
| Figure 4B |  |
| D2884 | MATa *cdc13-1 RAD53-3xHA::TRP1* |
| D3378 | MATa *cdc13-1 RAD53-3xHA::TRP1 cdc5-16::HIS3MX6* |
| D6788 | MATa *cdc13-1 RAD53-3xHA::TRP1 nud1-92::URA3MX6 cdc13-1 RAD53-3xHA::TRP1* |
| D9000 | MATa *cdc13-1 RAD53-3xHA::TRP1 spc110-7::kanMX6 spc72-8::kanMX6 nud1-92::URA3MX6* |
| Figure 4D |  |
| D6850 | MATa *RAD53-3xHA::TRP1* |
| D6853 | MATa *RAD53-3xHA::TRP1 nud1-92::URA3MX6* |
| D9086 | MATa *RAD53-3xHA::TRP1 spc110-7::kanMX6 spc72-8::kanMX6 nud1-92::URA3MX6* |
| Figure 5 |  |
| D6542 | MATa *cdc13-1 RAD53-3xHA::TRP1 ura3-1::3xGFP-CDC5::HIS3MX6::URA3 SPC29-RFP::HYGR* |
| D6808 | MATa *cdc13-1 RAD53-3xHA::TRP1 ura3-1::3xGFP-CDC5::HIS3MX6::URA3 SPC29-RFP::HYGR nud1-92::URA3MX6* |
| D8940 | MATa *cdc13-1 RAD53-3xHA::TRP1 ura3-1::3xGFP-CDC5::HIS3MX6::URA3 SPC29-RFP::HYGR spc110-7::kanMX6 spc72-8::kanMX6 nud1-92::URA3MX6* |
| Figure 6A |  |
| D8648 | MATa *CNM67-3xHA::URA3MX6 cdc15-2* |
| D8757 | MATa *CNM67-3xHA::URA3MX6 cdc5-99::HIS3MX6* |
| Figure 6C |  |
| D8648 | MATa *CNM67-3xHA::URA3MX6 cdc15-2* |
| D8650 | MATa *cnm67-16A-3xHA::TRP1 cdc15-2* |
| Figure 6D |  |
| D9440 | MATa *MPS3-3xHA::kanMX6 cdc15-2* |
| D9443 | *MATa MPS3-3xHA::kanMX6 cdc5-99::HIS3MX6* |
| Figure 6F |  |
| D9440 | *MATa MPS3-3xHA::kanMX6 cdc15-2* |
| D9446 | *MATa mps3-3A-3xHA::URA3MX6 cdc15-2* |
| Figure 7A |  |
| D4107 | *MATa* |
| D67 | *MATa nud1-18 (cdc18-1)* |
| D7034 | *MATa cnm67-16A::kanMX6* |
| D8065 | *MATa mps3-3A::kanMX6* |
| Figure 7B |  |
| D2884 | *MATa cdc13-1 RAD53-3xHA::TRP1* |
| D3378 | *MATa cdc13-1 RAD53-3xHA::TRP1 cdc5-16::HIS3MX6* |
| D7971 | *MATa cdc13-1 RAD53-3xHA::TRP1 cnm67-16A::kanMX6* |
| D8088 | *MATa cdc13-1 RAD53-3xHA::TRP1 mps3-3A::kanMX6* |
| Figure S1 |  |
| D6871 | *MATa hml::LEU2 hmr::URA3MX6 Pgal1-HO::URA3MX6* |
| D7205 | *MATa hml::LEU2 hmr::URA3MX6 Pgal1-HO::URA3MX6 nud1-28A::kanMX6* |
| D6904 | *MATa hml::LEU2 hmr::URA3MX6 Pgal1-HO::URA3MX6 nud1-18 (cdc18-1)* |
| D7122 | *MATa hml::LEU2 hmr::URA3MX6 Pgal1-HO::URA3MX6 nud1-92::kanMX6* |
| Figure S2 |  |
| D2884 | *MATa cdc13-1 RAD53-3xHA::TRP1* |
| D3378 | *MATa cdc13-1 RAD53-3xHA::TRP1 cdc5-16::HIS3MX6* |
| D9139 | *MATa cdc13-1 RAD53-3xHA::TRP1 spc110-7::kanMX6* |
| D9141 | *MATa cdc13-1 RAD53-3xHA::TRP1 spc72-8::kanMX6* |
| Figure S3A |  |
| D4107 | *MATa* |
| D67 | *MATa nud1-18 (cdc18-1)* |
| D8340 | *MATa spc42-Q110P::URA3MX6* |
| D6800 | *MATa nud1-28A::URA3MX6* |
| D6762 | *MATa nud1-92::URA3MX6* |
| Figure S3B |  |
| D2884 | *MATa cdc13-1 RAD53-3xHA::TRP1* |
| D3378 | *MATa cdc13-1 RAD53-3xHA::TRP1 cdc5-16::HIS3MX6* |
| D8834 | *MATa cdc13-1 RAD53-3xHA::TRP1 spc42-Q110P::URA3MX6* |
| Figure S3D |  |
| D6850 | *MATa RAD53-3xHA::TRP1* |
| D8800 | *MATa RAD53-3xHA::TRP1 spc42-Q110P::URA3MX6* |
| Figure S4A |  |
| D6871 | *MATa hml::LEU2 hmr::URA3MX6 Pgal1-HO::URA3MX6* |
| D7276 | *MATa hml::LEU2 hmr::URA3MX6 Pgal1-HO::URA3MX6 cnm67-16A::kanMX6* |
| Figure S4C |  |
| D6871 | *MATa hml::LEU2 hmr::URA3MX6 Pgal1-HO::URA3MX6* |
| D8097 | *MATa hml::LEU2 hmr::URA3MX6 Pgal1-HO::URA3MX6 mps3-3A::kanMX6* |

